# Targeting the homeodomain of ceramide-synthase can ameliorate insulin resistance

**DOI:** 10.1101/2025.07.24.666377

**Authors:** Christian Vera-Granda, Mariangela Sociale, Konstantin Beckschäfer, Ilja Kanevskij, Merlin Gerding, Franka Eckardt, Hannah Rindsfüsser, Christoph Thiele, Anna Ziegler, Margret H. Bülow, Reinhard Bauer

**Author notes:** Corresponding author: (RB).

## Abstract

Multiple studies have linked ceramide accumulation with insulin resistance and diabetes. Ceramide Synthases (CerS) are at the center of ceramide *de novo* formation. Impaired CerS activity leads to lower ceramide and resolves insulin resistance. Drosophila has only one CerS, named Schlank, which contains a catalytic lag1p motif and, like many CerS, a homeodomain regulating lipid homeostasis. How CerS homeodomains are associated with diabetes has been little studied. Here we demonstrate that, depending on the respective mutation in the CerS homeodomain high sugar diet (HSD)-induced insulin resistance is exacerbated or ameliorated. HSD shifts the profile of sphingolipids towards polyunsaturated longer sphingoid bases, systemic insulin signaling is reduced, as indicated by nuclear accumulation of FoxO, and secretion of insulin-like peptide 2 (DILP2) is impaired. Expression of a CerS variant with a mutation in the nuclear localization signal 2 within its homeodomain in the fat body improves systemic insulin signaling and DILP2 release. Thus, the CerS homeodomain may be a potential target to attenuate insulin resistance.

## Introduction

Increase of ceramides (Cer), which are intermediates in the biosynthetic pathway that produces complex sphingolipids (SL), is linked to obesity-associated metabolic dysfunction. Ceramide accumulation has been identified as a negative regulator of glucose tolerance and lipid metabolism (Raichur et al., 2014; Turpin et al., 2014) which leads to impaired energy homeostasis and eventually to insulin resistance (Chavez & Summers, 2012)

Ceramides are at the center of SL metabolism and are formed by the N-acylation of a sphingoid long-chain base by Ceramide synthases (CerSs) (Mullen et al., 2012). In mammals exist six CerS genes with different preferences for specific fatty acid chain lengths. For instance, CerS2 attaches very long fatty acyl CoAs such as C22–C24 to the sphingoid base and CerS5 and 6 have specificity for C14–C16. Recently, it was shown that Cer C16 produced by CerS6 play an essential role in the development of insulin resistance (Raichur et al., 2014; Turpin et al., 2014). In contrast, CerS6-deficient mice exhibited reduced Cer C16:0 and were protected from high-fat diet-induced obesity, resulting in amelioration of insulin resistance. Generally, interventions that prevent de novo Cer synthesis or lower Cer can resolve insulin resistance and prevent diabetes (Holland & Summers, 2008). However, whole body heterozygous deletion of Cers2 in mice drives a paradoxical compensatory increase in Cer C16:0, resulting in greater susceptibility to hepatic steatosis and diet-induced insulin resistance (Raichur et al., 2014). Therefore, CerSs are important targets for intervention, however, they may require careful assessment for side effects.

The *Drosophila melanogaster* genome encodes for only one CerS, named Schlank (Bauer et al., 2009). All CerS family members share a lag1p motif, which is evolutionarily conserved and required for CerS activity (Spassieva et al., 2006). Schlank is essential for the synthesis of a rather broad spectrum of Cer in *Drosophila* (Bauer et al., 2009; Ziegler et al., 2025). This is in contrast to the mammalian CerS family members, which use a specific subset of acyl-CoAs to produce Cer (Levy & Futerman, 2010). Like all vertebrate and insect CerS orthologues except CerS1 variants (Voelzmann & Bauer, 2010), the Drosophila CerS contains a homeodomain containing two nuclear localization sites (NLS1 and NLS2). All mouse and human homeodomain-containing CerSs also have two NLS motifs within the homeodomain, except for CerS3 (Voelzmann et al., 2016). We and others recently showed that the NLS2 site is required for DNA binding of Schlank to mediate transcriptional regulation (Sociale et al., 2018; Yuan et al., 2024). Mutation of nuclear localization site 2 (NLS2) within the homeodomain led to loss of DNA binding, deregulated gene expression, and NLS2 mutants can no longer adjust transcriptional response to changing lipid levels. Now, a recent study showed that a glutamine-to-alanine substitution at position 115 (E115A) in the homeodomain of CerS2, just upstream of the NLS2, is associated with enhanced diet-induced glucose intolerance and hepatic steatosis (Heinitz et al., 2024; Nicholson et al., 2021).

Here, we aimed to further analyse how different mutations within the homeodomain affect DNA binding, SL profiles, and metabolic outcomes e.g. insulin resistance. We wanted to find out whether and how the lag1p motif and homeodomain come into play with different degrees of emphasis during the development of insulin resistance using a type 2 diabetes (T2D) model generated by consuming a high-sugar diet (HSD).

## Results

### Phenotypic analysis of CerS Schlank Homeodomain Mutants

We have recently established an animal model that allows us to study domain-specific functions of the Drosophila CerS homeodomain and of the catalytic lag1p motif *in vivo* (Fig 1A) (Sociale et al., 2018). Mutation of nuclear localization site 2 (NLS2) within the homeodomain leads to loss of DNA binding, deregulated gene expression, and no longer adjusts transcriptional response to changing lipid levels. To further analyse the effects of manipulations within the homeodomain, we now targeted the NLS1 site and introduced a glutamine-to-alanine substitution at position 118 (E118A) corresponding to the single-nucleotide polymorphism (human SNP; rs267738 or CerS2 E115A) detected in the homeodomain of human CerS2. The substitution at E115A within CERS2 is a strong candidate linking SL metabolism with defects in glucose homeostasis of mammalian CerS2 (Khan et al., 2025; Nicholson et al., 2021). The mutant variants were reintegrated into the native *schlank* locus in the *schlank^KO^ ^founder^ ^line^*, resulting in knock-in (KI) KINLS1 and KIE118A fly lines, just like we had done before to generate the KINLS2 mutant variant (S1 Fig). As a control, we used KIWT animals, which are indistinguishable from wild-type (WT) flies (Sociale et al., 2018).

**Fig 1.**
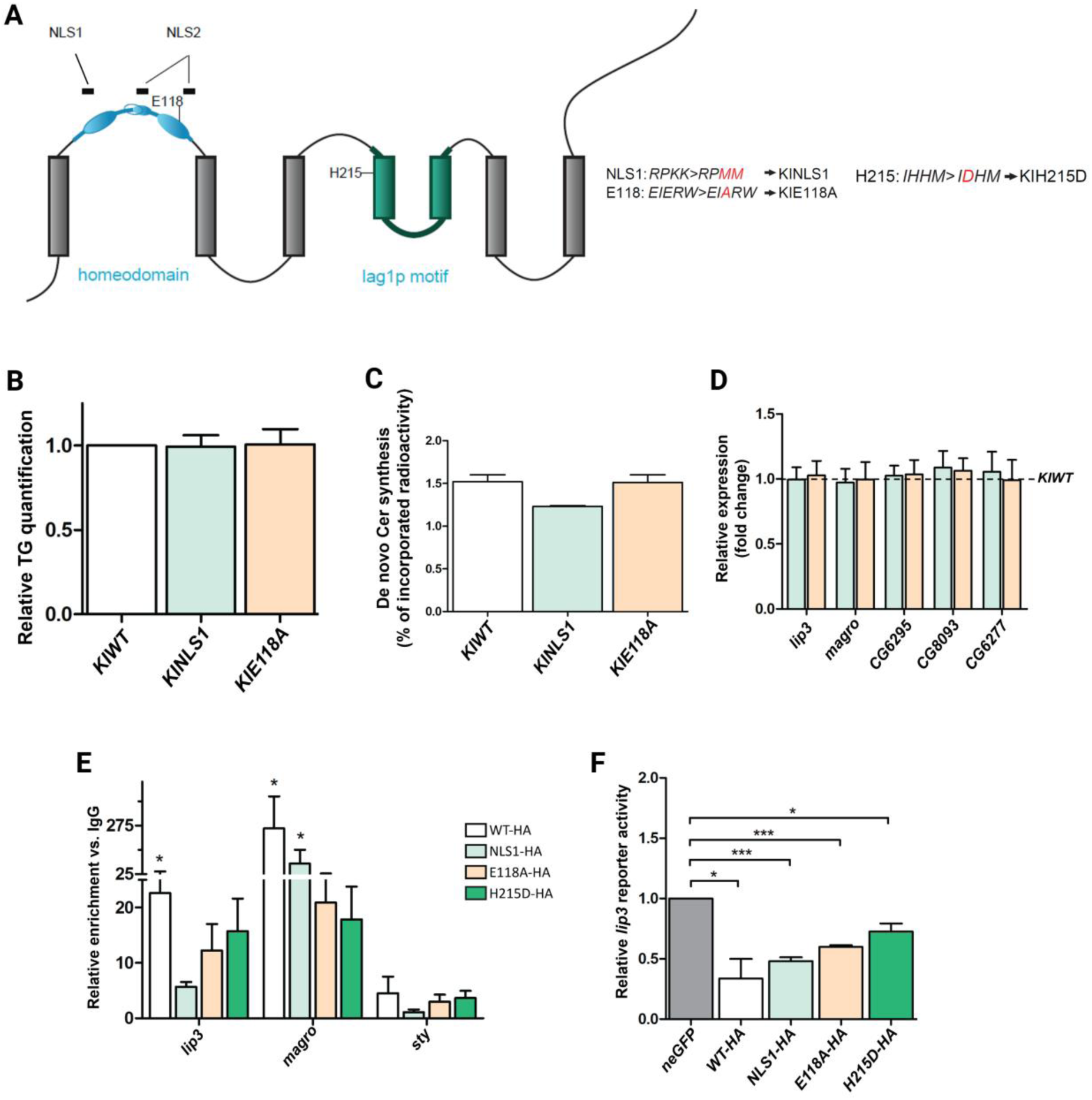
Characterization of Drosophila CerS homeodomain mutants. (A) schematic representation of CerS topology (modified after Thidar et al., 2018). CerS Schlank protein topology (modified according to Sociale et al., 2018). Grey boxes indicate transmembrane domains (TM), blue marks the homeodomain and its helices, and green the Lag1p motif. Black boxes above the homeodomain reveal the Monopartite-like NLS1 (RPKK, aa78–81) and an atypical bipartite-like NLS2 (RLDKKK-X19RLRR, aa97–125). (Á) Mutations of newly generated (KINLS1, KIE118A) and of already existing (KINLS2, KIH215D) CerS mutant alleles: KINLS1 two lysines (K) are changed into methionine (M) RPKK>RPMM; KIE118A glutamate changed to alanine E>A; KINLS2 two arginines (R) are changed into alanine (RLRR-> ALAR); KIH215D histidine changed into aspartate H>D. (B) Total body fat (TG) in KINLS1 and KIE118A L3 larvae is similar to that of the control. Relative TG content was normalized to dry weight. (C) [1-^14^C]-acetic acid was used to perform a metabolic labelling experiment in complete larvae to determine d*e novo* Cer generation. Equal amounts of radioactivity were applied to TLC plates and % of the incorporated label was quantified. Biosynthesis of Cer in KIE118A mutant larvae showed no difference in comparison to KIWT controls whereas KINLS1 shows a moderate reduced activity. (D) Quantification of transcript levels of tested lipases in KINLS1 and KIE118 L3 larvae was done by RT–qPCR (KIWT was used as control). Error bars indicate ±SEM; * p<0.05, ** p<0.01, *** p<0.001. (E) NLS1 and E118A homeodomain mutant variants bind *lip3* and *magro* promoter regions. Quantification by RT–qPCR of ChiP material from S2 cell extract transfected with WT-HA, NLS1-HA, E118A-HA, and H215D-HA (n=3), using α-HA antibody. Promoter regions assayed were those of *lip3*, *magro* and *sty*, (negative control). Expression was normalized to the relative expression of the IgG sample Error bars indicate ±SEM. (F) Relative luciferase induction upon expression of either WT-HA, NLS1-HA, E118A-HA, and H215D-HA is indicated, normalized to renilla luciferase control transfection. The control bar is in reference to luciferase activity upon UAS-GFP control expression (n = 3). Error bars indicate ±SEM.

KIE118A and KINLS1 mutant animals are phenotypically similar to wild-type animals, except for a moderate delay in KIE118A lines reaching the pupariation stage (S2A Fig). There were also no differences in triacylglycerides (TG), *de novo* Cer synthesis and expression levels of lipases (Fig 1B-D), among which lip3 and magro are direct Schlank targets.

This is a striking contrast to the KINLS2 mutants, for which we showed reduced TG levels and chronically upregulated lipase expression. The negative regulatory effect on lipase transcription by directly binding to their promoters is lost in the NLS2 variant since it has lost the ability to bind DNA (Sociale et al., 2018; Yuan et al., 2024). Therefore, we wanted to know whether the KINLS1 and KIE118A mutated variants can bind DNA by performing chromatin immunoprecipitation (ChIP) experiments.

To this end, we used the following Schlank variants fused with a hemagglutinin tag (HA-tag): wild-type (HA-WT), the homeodomain NLS1 and E118A (NLS1-HA, E118A-HA) mutants, and the Cer synthesis-deficient H215D-HA variant, which has a mutation in the catalytic lag1p motif. After expression of these variants in S2 cells, we performed ChIP. As negative controls, we performed immunoprecipitations (IPs) using anti-IgG antibodies. The catalytically inactive H215D-HA variant for which DNA binding capacity was already shown (Sociale et al., 2018) was used as a positive control. RT-qPCR on the immunoprecipitated material revealed that promoter regions of lip3 and magro were enriched in the HA-ChIP from all extracts compared with the genomic region of an unrelated genomic region of sprouty (sty), which was used as a negative control. In accordance with ChIP results, the expression of either wild-type, NLS1-HA, E118A-HA, H215D-HA variants, all binding DNA, could repress *lip3* luciferase activity (Fig 1E-F). This is in agreement with previous experiments, which revealed a repression of lip3 transcription only upon expression of CerS variants where DNA-binding capability was not affected (Sociale et al., 2018).

In sum, these data demonstrate that all CerS mutated variants analysed so far, with the exception of NLS2 mutants, bind DNA, did not impact TG- or expression-levels of lipases and had no particular phenotypic abnormalities.

### Drosophila CerS E118A and NLS1 homeodomain mutants increase the predisposition to insulin resistance upon an HSD

Previously, it was shown that mice heterozygous for CerS2 or homozygous knock-in CerS2 E115A mice were indistinguishable from wild-type controls when fed a chow diet but were predisposed to diet-induced glucose intolerance when fed a high-fat diet (HFD) (Nicholson et al., 2021; Raichur et al., 2014). The insulin pathway is highly conserved from mammals to *Drosophila* and plays fundamentally the same physiological functions (Dutriaux et al., 2013; Teleman, 2010). Drosophila produces eight insulin-like peptides (ILP) called DILPs, which have structural and functional similarities with vertebrate insulin-like growth factor and insulin. Three of the eight DILPs (DILP 2, 3 and 5) are produced in the brain’s insulin-producing cells (IPCs), which are homologous to pancreatic beta cells (Géminard et al., 2009; Rulifson et al., 2002) and bind upon release from the IPCs to the insulin receptor in peripheral tissues to promote growth and nutrient utilization (P et al., 2020; Pasco & Léopold, 2012; Semaniuk et al., 2021).

Therefore, we decided to raise the KINLS1 and KIE118A animals on HSD (Musselman et al., 2011, 2019; Na et al., 2013). The CerS homeodomain mutated larvae, as well as the wildtype control reared on HSD, were reduced in size and weight, had elevated hemolymph glucose levels, and showed a strong delay in larval developmental rate. This phenotype was even more pronounced in the KIE118A and KINLS1 animals (Fig 2A-D), indicating characteristic signs highly associated with insulin resistance (Azmin et al., 2025; Meshrif et al., 2022; Musselman et al., 2011; P et al., 2020).

**Fig 2.**
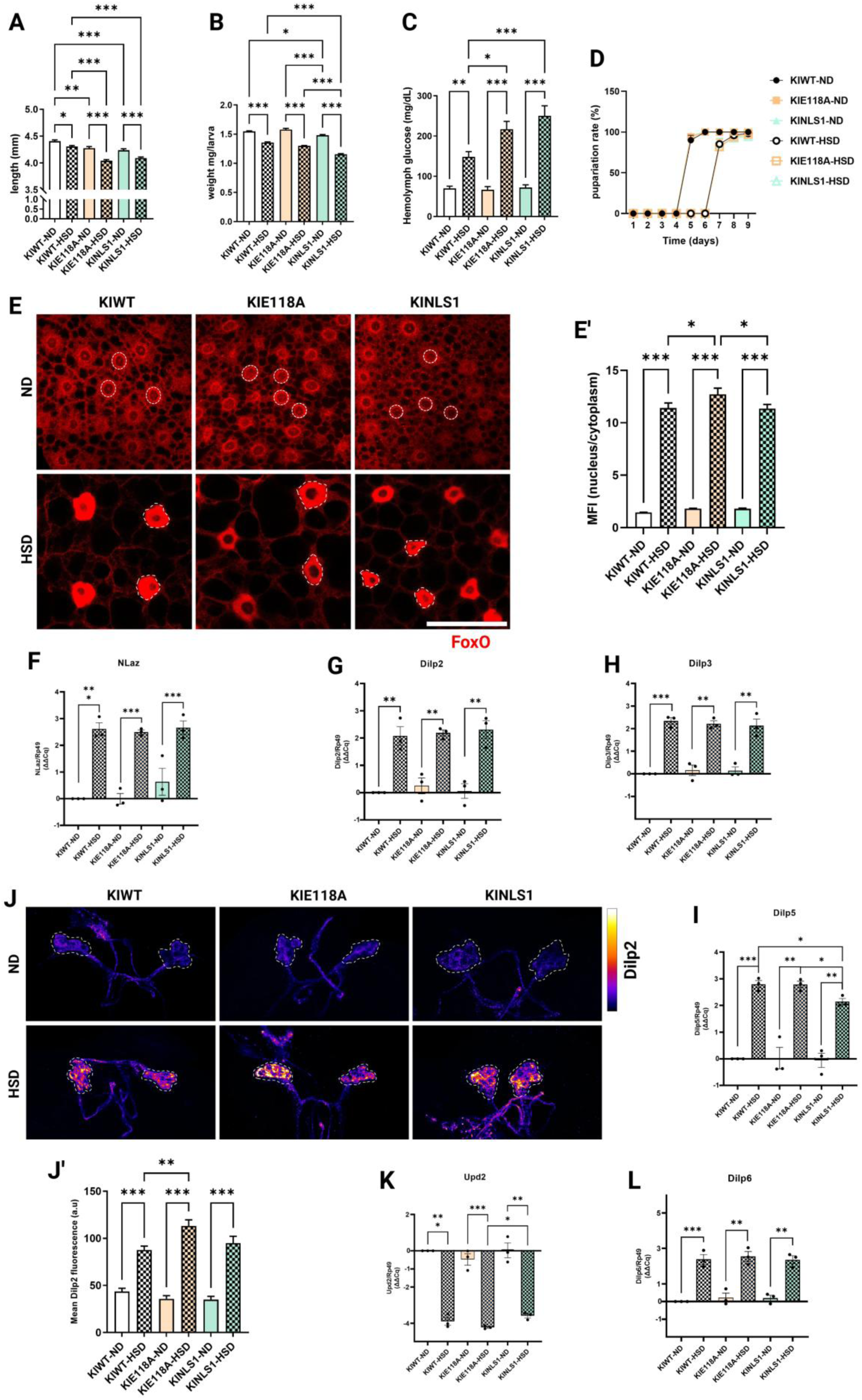
Drosophila CerS E118A and NLS1 homeodomain mutants promote metabolic disturbance upon an HSD. (A-B) Lengths and body weights of L3-instar larvae show a significant decrease in KINLS1 and KIE118A compared to KIWT when exposed to 1M HSD and 0.15M ND ((A): n=60; (B): n≥15). (C) Hemolymph glucose concentrations of L3-instar mutated larvae increase significantly when reared HSD (n=30). (D) Developmental time frame to pupariation of L3 larvae shows 2-3 days delay when animals are fed HSD compared to ND (n=3). (E) Immunostaining of FoxO in L3-instar larvae FB of CerS homeodomain mutated variants determining their nuclear/cytoplasmic localization of FoxO upon HSD and ND. The dashed outlines show the area of the nuclei. Scale bar = 50µm. (E’) Relative mean fluorescence intensity (MFI) of FoxO staining (nucleus/cytoplasm) in FB cells of KINLS1 and KIE118A homeodomain mutated larvae reared in HSD or ND, showing increased nuclear localization of FoxO in KIE118A mutants upon HSD (n≥15). (F-I) Quantification of the dilps transcript levels by RT-qPCR of CerS homeodomain mutated variants compared with the wild-type control in the FB (F) nlaz; in the brain (G) dilp2, (H) dilp3 and (I) dilp5 transcript levels. There is a consistent and significant upregulation of these mRNA transcripts in the FB and larval brains of animals reared HSD (n=3 biological replicates). (J) Immunostaining determining the accumulation of DILP2 peptide through the fluorescence intensity of an α-DILP2 antibody in IPCs of L3-instar mutated larvae and WT (dashed outline), showing increased DILP2 accumulation upon a prolonged HSD consumption. (J’) Quantification of relative mean DILP2 fluorescence intensity showing its significantly increased accumulation in IPCs of KIE118A mutated larvae as compared to control IPCs of animals fed HSD (n≥15). (K-L) Quantification of the transcript levels of (K) upd2, and (L) dilp6 by RT-qPCR in the FB cells of CerS homeodomain mutated variants and the wild-type control. The dilp6 transcript levels are significantly upregulated while the upd2 transcript levels are downregulated in animals reared HSD (n=3 biological replicates). Expression level normalized on rp-49, and the ΔΔCq is represented and used in statistical analysis. Error bars from each graph mentioned above indicate ±SEM. Unpaired two-tailed t-test and one-way ANOVA followed by Tukey’s multiple comparisons tests were used to derive P-values * p<0.05, ** p<0.01, *** p<0.001.

Now we sought to determine whether these homeodomain mutants have an influence on the development of insulin resistance induced by HSD. Hence, we examined key markers for activity of insulin signaling or insulin resistance, the insulin-dependent forkhead transcription factor (FoxO) and Lipocalin-encoding Neural Lazarillo (Nlaz), an insulin resistance marker.

When insulin signaling is active, FoxO is phosphorylated and translocated to the cytoplasm. In contrast, when insulin signaling is not active or in a state of reduced sensitivity to insulin, FoxO is relocated to the nucleus (Lee & Dong, 2017; Ni et al., 2007; P et al., 2020). Using a FoxO antibody we detected FoxO protein primarily in the cytoplasm in the larval fat body (FB; liver equivalent) on ND. However, it was considerably nuclear localized on HSD, being markedly more nuclear in the KIE118A, while in the KINLS1 it was similar to the KIWT (Fig 2E, E’). Furthermore, we detected that the expression of insulin resistance marker nlaz (Lourido et al., 2021; Pasco & Léopold, 2012), was upregulated in FB cells of CerS homeodomain mutants, as well as in the wildtype control on HSD (Fig 2F), confirming the decrease in insulin sensitivity. This evidence ratifies that chronic HSD consumption leads to insulin resistance, but also, the increased nuclear localization of FoxO in the KIE118A mutants after HSD might suggest a potential predisposition to the onset of insulin resistance in the periphery.

Next, we explored whether DILP2 production and release were impaired. We focused our analysis on DILP2, for which we can easily follow using a specific antibody. While an acute starvation leads to a strong accumulation of DILP2, transferring starving animals back on amino acid-rich food reverses this accumulation (Géminard et al., 2009). For that reason, we first tested whether the release of DILP2 in the IPCs was affected by CerS homeodomain mutants in starving and refeeding conditions. As expected, using a specific antibody against DILP2, we observed that the DILP2-containing secretion granules tend to accumulate slightly more in the IPCs of KIE118A under acute starvation, but also after refeeding conditions (yeast food) for up to 1.5 h in comparison to KINLS1 and wild-type control (S4A, A’ Fig). Considering that the exacerbated production and disturbed secretion of DILPs, including hyperglycemia, is characteristic of peripheral insulin resistance, we hypothesized that the CerS homeodomain mutants most likely impact the expression and release of DILPs from IPCs under an HSD regime.

To test this, we first measured mRNA levels of dilps, which are produced in the IPCs, dilp2, 3 and 5, following long-term high-sugar consumption. As expected, dilp levels were upregulated, but we were unable to detect a significant difference between CerS homeodomain mutants and control (Fig 2G-I). Therefore, we next tested for the DILP release from IPCs after a high-sugar consumption. Indeed, we found that under this condition (P et al., 2020; Pasco & Léopold, 2012), the accumulation of DILP2 in the IPCs was considerably increased. Strikingly, the amount of DILP2 in the IPCs of KIE118A mutants was significantly higher than in the control, while DILP2 increase in IPCs of KINLS1 mutants was comparable to the control (Fig 2J, J’). Our results for KIE118A resemble the glucose intolerance and impaired insulin secretion observed in CerS2 E115A knock-in mice (Heinitz et al., 2024; Khan et al., 2025). In sum, the E118A mutation within the homeodomain had no impact on dilp-expression, but DILP-release to the systemic circulation is apparently affected.

In accordance with the strong nuclear localization of FoxO, these data suggest that the impact of the E118A mutant on the insulin sensitivity is not only restricted to the FB cells but also affects the DILP secretion from the IPCsIn order to know whether insulin release-related factors produced in the FB could be involved in this process, we determined the expression of upd2 and dilp6 in the FB. Upd2 is a cytokine that promotes the release of DILP2 by IPCs, which is upregulated as an acute response to HSD decaying at 72h after egg laying (AEL) (Ingaramo et al., 2020; Lourido et al., 2021). Conversely, the DILP6 promotes the inhibition of DILP2 release from IPCs to the systemic circulation (Bai et al., 2012; Lourido et al., 2021). In line with the observed strong accumulation of DILP2 in the IPCs after HSD consumption, mRNA levels of upd2 were decreased and expression of dilp6 increased in FB of L3 instar larvae (72-96h AEL) (Fig 2K, L). However, no noticeable difference was observed between the homeodomain mutants and the controls.

Nonetheless, the increased accumulation of DILP2 in IPCs and the marked nuclear FoxO localization in KIE118A mutants still point to an increased potential to disrupt insulin signaling on HSD as compared to wildtype animals. This may not be reflected at the transcriptional level but possibly through changes in body fat metabolism (Bauer et al., 2009; Voelzmann et al., 2016) or lipid composition.

### CerS homeodomain mutations alter the lipid profile upon an HSD

To examine the effect of Drosophila CerS homeodomain mutants on body fat upon caloric overload (HSD), we analysed triacylglycerides (TG) in lipid droplets and the expression of TG metabolism-related genes.

To this end, we first analysed the content and distribution of TG in lipid droplets by staining the neutral lipids with Nile red in the FB, the major lipid storage organ. We observed fat accumulation and an increasing lipid droplet area when reared on HSD (Fig 3A-A’) for the CerS homeodomain mutants and wildtype control. Consistent with this and the increased total TG levels in the whole body and the FB (Fig 3B, C) lipid storage droplet-1 (lsd-1) and phosphoenolpyruvate carboxykinase (pepck) expression are upregulated (Fig 3D, E) whereas mRNA levels of lipases cg8093 (Vaha), cg6277, and lip3 (Fig 3F-H) in the whole larval body are downregulated. For Vaha it was furthermore found that it stimulates IPCs to release DILPs into the circulatory system, which is severely reduced after its loss (Singh et al., 2024). However, there was no particular difference between mutants and controls in body fat content and the regulation of genes involved in lipolysis and lipogenesis between CerS homeodomain mutants and wild-type animals.

**Fig 3.**
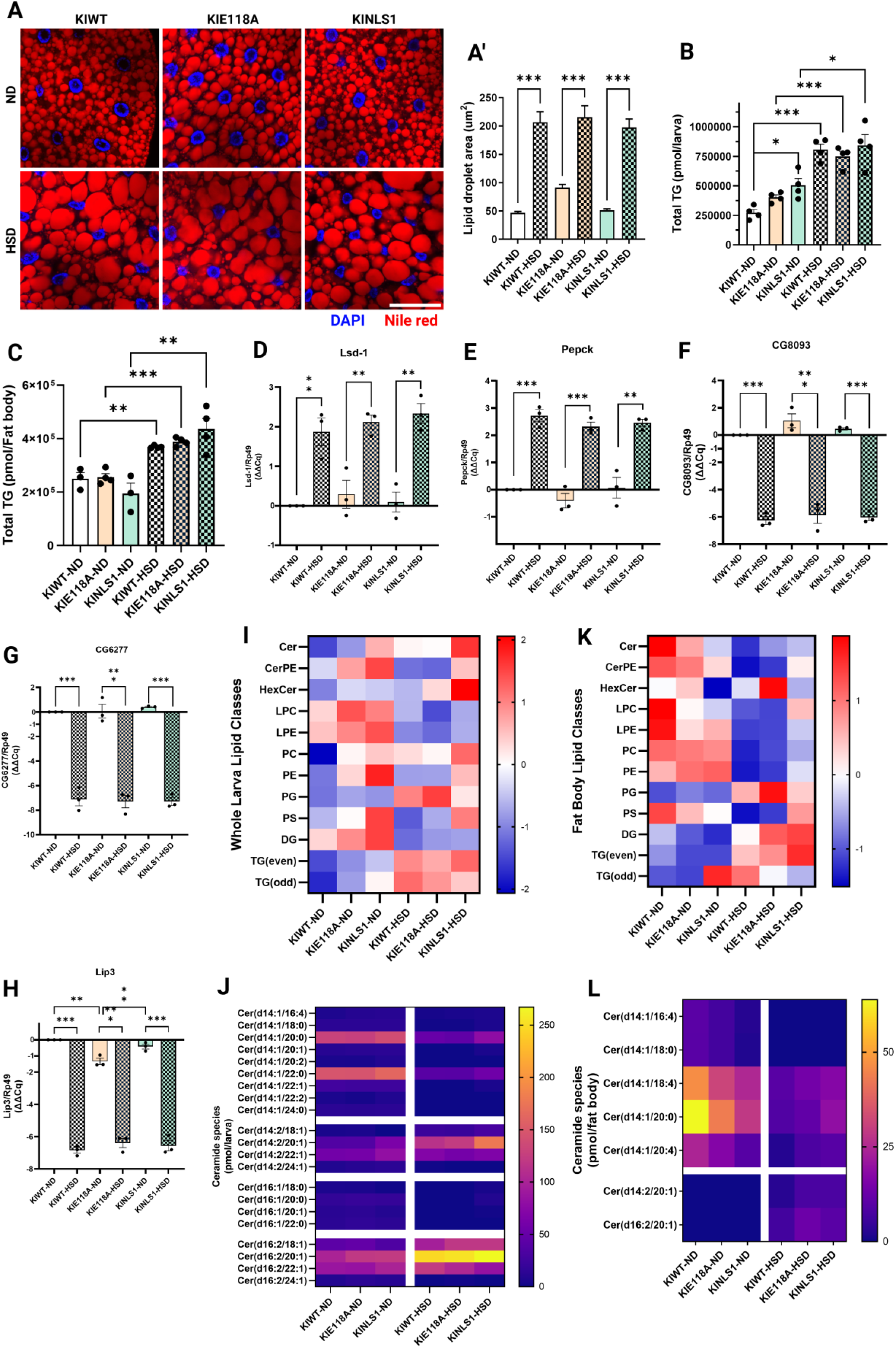
Drosophila CerS Schlank homeodomain mutations alter the lipid profile upon an HSD. (A) Neutral lipid (Nile Red) staining of FB cells from KINLS1 and KIE118A homeodomain mutated L3-instar larvae reared HSD or ND compared with control wildtype. (A’) Quantification in two-dimensional confocal slices of lipid droplet area (n≥15). The homeodomain-mutated L3 larvae upon HSD show a high increase in lipid droplet area compared to the animals reared in ND. (B) Total TGs extracted from the whole larval body upon ND or HSD and measured by tandem shotgun MS, indicating an upregulation of total TGs when animals are reared HSD (n=4 biological replicates). (C) Total TGs extracted from FB of mutated and wild-typic animals upon ND or HSD and measured by tandem shotgun MS, indicating an upregulation of total TGs when animals are reared HSD (n=4 biological replicates). (D-E) Relative expression by qPCR of lsd-1 and pepck. It shows the significant upregulation of (D) lsd-1 and (E) pepck in FB cells from larvae-reared HSD compared to larvae-reared ND (n=3). (F-H) Relative expression by RT-qPCR of lipases (F) cg8093, (G) cg6277, and (H) lip3 in the whole larval body of CerS homeodomain mutated variants compared to the wildtype larvae. The transcript levels of lipases are significantly downregulated in animals when reared on HSD (n=3). Expression level normalized on rp-49, and the ΔΔCq is represented and used in statistical analysis. (I) Heatmap summarizing the representative lipid classes quantified relative to lipid standards in each replicate of one larval body per group and standardized into z-score from whole L3-instar mutated and wildtype larvae reared on ND and HSD. Lipids were extracted and measured by tandem shotgun MS (n=4 biological replicates). (J) Heatmap representation of ceramide species extracted from whole L3-instar mutated and wildtype larvae reared on ND and HSD and measured by tandem shotgun MS (n=4 biological replicates). (K) Heatmap summarizing the representative lipid classes quantified relative to lipid standards in each replicate of one FB per group and standardized into z-score from CerS homeodomain mutated and wildtype FB of animals reared on ND and HSD. Lipids were extracted and measured by tandem shotgun MS (n=4 biological replicates). (L) Heatmap of ceramide species extracted from homeodomain mutated and wildtype FB cells of animals reared on ND and HSD and measured by tandem shotgun MS (n=4 biological replicates). Abbreviations: Ceramide (Cer), Ceramide phosphatidylethanolamine (CerPE), Hexosylceramide (HexCer), Diacylglyceride (DAG), Lyso-phosphatidylcholine (LPC), Lyso-phosphatidylethanolamine (LPE), Phosphatidylcholine (PC), Phosphatidylethanolamine (PE), Phosphatidylglycerol (PG), Phosphatidylserine (PS). Error bars are ± SEM. Unpaired two-tailed t-test, one-way and two-way ANOVA followed by Tukey’s multiple comparisons tests were used to derive P-values; * p<0.05, ** p<0.01, *** p<0.001.

Next, to better understand the impact of CerS E118A and NLS1 homeodomain mutations on the lipid composition we evaluated the lipidome of complete larvae and FB, the primary lipid storage and metabolic tissue by shotgun tandem mass spectrometry (MS). We observed that several lipid classes were increased in the whole larval body of CerS homeodomain mutants compared to wildtype control on ND. Among the lipid classes that were increased, we found SL like Cer (mainly in KINLS1) and ceramide phosphoethanolamine (CerPE); neutral lipids like diacylglycerides (DG); and phospholipids (PL) like lysophosphatidylcholine (LPC), lysophosphatidylethanolamine (LPE), phosphatidylcholine (PC) phosphatidylethanolamine (PE) and phosphatidylserine (PS) (Fig 3I). In contrast, we detected upon feeding HSD an increase in the Cer and hexosylceramide (HexCer) contents, mainly in KINLS1 (Cer ∼1182,98 pmol/larva, and HexCer ∼1805,35 pmol/larva in average) and not quite as much in KIE118A (Cer ∼1033,97pmol/larva, HexCer ∼1232,58 pmol/larva in average), in comparison to the control (Cer ∼1038,37 pmol/larva, HexCer ∼944,52 pmol/larva) (Fig 3I). We also observed that the content of phosphatidylglycerol (PG) was increased in animals after HSD-feeding, standing out mainly in KIE118A mutants, suggesting a response to an intrinsic inflammation likely caused by an obesogenic diet. PGs might be incorporated into cardiolipins, improving mitochondrial activity and inhibiting inflammation (Chen et al., 2018; Chu et al., 2022; Schleh et al., 2023). Furthermore, we noted that in both Cer and HexCer species of mutant and control larvae reared on HSD, the number of sphingoid bases with two double bonds, such as d14:2/20:1, d16:2/18:1, and d16:2/20:1, was markedly increased. In contrast, Cer species containing monounsaturated sphingoid bases such as Cer (d14:1/20:0) and Cer (d14:1/22:0) were decreased (Fig 3J; S4 E-E’, I-I’ Fig). It is also interesting to note that the latter are the most abundant Cer under ND conditions (Bauer et al., 2009; Ziegler et al., 2025).

In contrast to the whole larval body lipidome, the FB lipidome presented reduced SL, such as Cer and CerPE, except for HexCer, which is increased predominantly in KIE118A mutants (Cer ∼53,86 pmol/larva, CerPE ∼66,13 pmol/larva, and HexCer ∼311,91 pmol/larva on average) upon HSD (Fig 3K). The reduced FB content of Cer in the KIE118A and KINLS1 mutants under an ND regime is consistent with a potential loss in CERS2 activity that was observed for the human SNP (E115A) variant in liver homogenate (Nicholson et al., 2021). However, under an HSD regime, the Cer content in the KIE118A and KINLS1 mutants was not significantly reduced compared to wildtype control. Among the most remarkable decreased Cer species in FB was again Cer (d14:1/20:0), which was also reduced in the whole larval body when reared on HSD and which is one of the main Cer species generated *de novo* by CerS Schlank. In contrast, as observed in the whole larval body, we detected an increase in Cer (d14:2/20:1) and Cer (d16:2/20:1), which were surprisingly just detectable upon an HSD, but not in ND (Fig 3L; S4D Fig) and more pronounced in the NLS1 and E118A variants.

In mammals, obesity- or insulin resistance-related alterations are mainly attributed to specific N-acyl-chain length Cer (Turpin et al., 2014; Turpin-Nolan et al., 2019). A potential insulin resistance-related Cer species in Drosophila is not yet known. However, it is noteworthy that we detected a changing trend towards doubly unsaturated sphingoid bases in the lipid profile in our HSD T2D model. These changes in lipid species may have consequences for membrane lipid composition and function, leading to insulin-related metabolic disease.

### CerS NLS2 homeodomain and H215D catalytic lag1p motif mutations alleviate insulin resistance

Next, we asked what influence the mutation of the NLS2 motif within the homeodomain would have on the development of insulin resistance and the lipid profile after HSD feeding. We previously showed that NLS2 mutation leads to loss of DNA binding and deregulated gene expression. Moreover, KINLS2 mutants have the strongest phenotypes of all the homeodomain mutants we generated. They are severely delayed in development and growth, with many dying during all larval stages, and only about 10% of the initial population ecloses as small-sized adults of normal proportion with locomotion deficits (Sociale et al., 2018). In addition, we decided to analyse the enzymatically inactive H215D CerS variant (Sociale et al., 2018; Ziegler et al., 2025) with a mutation within the catalytical lag1p motif. These animals show lethality in the larval first instar stage, dying even earlier than KINLS2 animals.

For this reason, we could not analyse these mutant variants when ubiquitously expressed as we had done for the KINLS1 and KIE118A variants and chose to conditionally express the H215D and NLS2 CerS variants in the FB, which is functionally equivalent to mammalian adipocytes and liver (Zheng et al., 2016). To this end, we generated so-called allele switch lines switching from a wild-type *CerS Schlank* flanked by FRT sites to a wild-type HA-tagged *CerS Schlank* (as control) or to either the HA-tagged H215D- or the HA-tagged NLS2-variant, when the Flippase (Flp) is present. Here we used the r4-GAL4 driver line to specifically express UAS-Flippase in the larval FB, leading to the excision of the wild-type *CerS Schlank* and switching to the mutated variant (Fig 4A-A’; S3 Fig).

**Fig 4.**
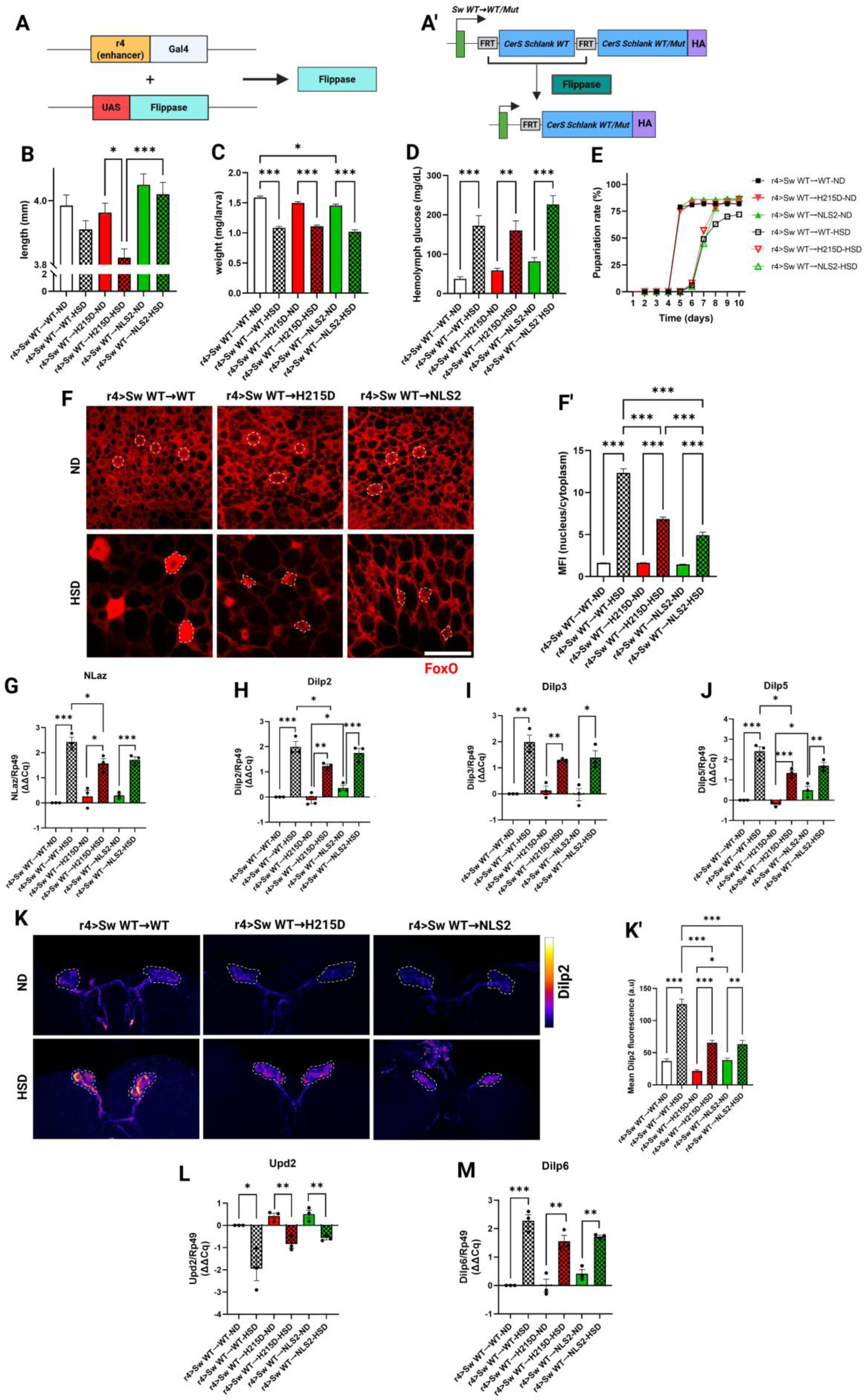
Drosophila CerS NLS2 homeodomain and H215D catalytic domain mutations in FB alleviate HSD diet metabolic disturbance. (A-A’) The expression of the CerS Schlank mutants and wild-type control in the FB was obtained by crossing Sw WT→H215D (catalytic domain), Sw WT→NLS2 (homeodomain), and Sw WT→WT (wildtype) with the r4-Gal4 driver. The allele switch is induced by excision of *Schlank WT* by a Flippase (Flp). Subsequently, the HA-tagged *Schlank WT* as a control or the HA-tagged *Schlank Mut* allele is expressed. Abbreviations: FRT: flippase recognition targets, HA: hemagglutinin tag, UAS: upstream activation sequence. (B) Lengths, (C) body weights, and (D) hemolymph of L3-instar CerS FB mutated and wildtype larvae are significantly affected when reared in 1M HSD compared to 0.15M ND. While length and body weight are decreased, the hemolymph glucose is increased upon an HSD ((B): n=50; (C): n≥15); (D): n=16). (E) Developmental time frame to pupariation of larvae bearing CerS mutations in FB. It shows a 2-3 days delay when animals are fed HSD compared to 3-5 days in wildtype control (n=3 biological replicates). (F) Immunostaining of FoxO in L3-instar larvae FB of catalytic domain and homeodomain mutated variants compared to control larvae, determining their nuclear/cytoplasmic localization of FoxO upon HSD and ND. The dashed outlines show the area of the nuclei. Scale bar = 50µm. (F’) Quantification of the relative mean fluorescence intensity (MFI) of FoxO staining (nucleus/cytoplasm) shown in (F), showing less nuclear localization of FoxO in the CerS mutated variants in comparison to wild-type control upon HSD (n≥15). (G-J) Quantification of the dilps transcript levels by RT-qPCR in the FB (G) nlaz and the brain of larvae (H) dilp2, (I) dilp3, and (J) dilp5 of CerS catalytic domain and homeodomain mutants in the FB. The nlaz transcript levels are less upregulated in CerS-mutated larvae compared to the control. A significant upregulation of the mRNA transcripts is shown in the larval brains of animals reared on HSD (n=3 biological replicates). (K) Immunostaining determining the accumulation of DILP2 peptide through the fluorescence intensity of an α-DILP2 antibody in IPCs of larvae expressing CerS mutations in the FB (dashed outline), showing increased Dilp2 accumulation upon prolonged HSD consumption. (K’) Quantification of relative mean DILP2 fluorescence intensity presented in (K), showing that DILP2 peptide was significantly less accumulated in IPCs of larvae bearing CerS mutations in FB, as compared to control IPCs of animals fed HSD (n≥15). (L-M) Quantification of the transcript levels of (L) upd2, and (M) dilp6 by RT-qPCR in the FB. The dilp6 transcript levels are upregulated upon HSD. In contrast, the upd2 transcript levels are less downregulated in FB of CerS-mutated larvae reared on HSD compared to the control (n=3 biological replicates). Expression level normalized on rp-49, and the ΔΔCq is represented and used in statistical analysis. Error bars from each graph mentioned above indicate ±SEM. Unpaired two-tailed t-test and one-way ANOVA followed by Tukey’s multiple comparisons tests were used to derive P-values * p<0.05, ** p<0.01, *** p<0.001.

Control, H215D- or the NLS2 animals were raised HSD, and as expected, a long-term consumption of HSD led to a considerably reduced size and weight, and hemolymph hyperglycemia. We did not observe a significant difference for all these characteristics associated with insulin resistance when wildtype or the mutant variants were expressed in the FB (Fig 4B-D). However, interestingly, the developmental rate was noticeably improved in H215D and NLS2 mutants, increasing the percentage of pupariation rate and, consequently, their survival on HSD (Fig 4E). The improvement in the developmental rate suggests that the mutation in the CerS catalytic domain or of the NLS2 site in the homeodomain provides an advantage over the wild-type control under a caloric overload.

To better characterize the potential attenuation effect of the H215D mutation in the CerS catalytic domain and its NLS2 mutation in the homeodomain on insulin resistance, we analysed the localization of FoxO as a biomarker of metabolic imbalance and a readout for insulin sensitivity in these mutants reared on HSD and ND (P et al., 2020; Zang et al., 2024). Larvae, which were fed on ND did not display significant differences in their cytoplasmic and nuclear localization. Noteworthy, we found strikingly less nuclear localization of FoxO in the FB of the mutant animals when reared on HSD. Surprisingly, the NLS2 mutants even showed significantly less nuclear FoxO localization than the catalytic inactive H215D variant (Fig 4F-F’). Besides, we assessed the expression of the insulin resistance marker (nlaz), in response to HSD (Lourido et al., 2021; Pasco & Léopold, 2012). Although the expression of nlaz was upregulated in our CerS mutants upon HSD, it was significantly lower expressed in the H215D variant and not significantly, but still lower expressed in NLS2 variant as compared to wildtype control (Fig 4G).

Since the FB-expressed Drosophila CerS catalytic domain and homeodomain NLS2 mutant variants had an improving impact on modulating insulin signaling-related factors FoxO and nlaz, we asked whether the release of DILP2 from the IPCs might also be improved. As done for KIE118A and KINLS1 mutants, we tested whether the release of DILP2 in the IPCs was affected in starving and refeeding conditions (S5A, A’ Fig) when the NLS2 or H215D variant was expressed in FB. Unlike the increased accumulation of DILP2 in the IPCs of the KIE118A mutant after starvation and refeeding (S4A, A’ Fig), the accumulation of DILP2 in NLS2 and H215D mutants was comparable to the wild-type control under these conditions (S5A, A’ Fig). Next, we asked whether dilp mRNA expression and DILP secretion are affected in these mutants under long-term high-sugar consumption.

Thus, we measured the expression levels of DILP2, 3 and 5 produced in the IPCS, in dissected brains. mRNA levels of these dilps between CerS mutant variants and the control were comparable on ND. On HSD, we noted a significant increment in their expression in line with the results of previous studies (Lourido et al., 2021; Musselman et al., 2011; Pasco & Léopold, 2012). Noteworthy, a tendency towards less strong upregulation of dilps was observed in brains of CerS mutants (Fig 4 H-J).

Therefore, we hypothesized a reduced accumulation on HSD as well. Indeed, by using an anti-DILP2 antibody, we detected a significant reduction in DILP2 accumulation in IPCs of the FB-expressed NLS2 or H215D mutants (Fig 4K-K’), suggesting an improvement in insulin secretory granule turnover. These data also suggest that a potential crosstalk mediated by organ-to-organ signals and modulated by the Drosophila CerS catalytic domain and homeodomain might exist between FB and IPCs.

For this reason, we checked now for the expression of upd2 and dilp6 in the FB as we had done before for the KIE118A and KINLS1 mutants. While Upd2 promotes the release of DILP2, DILP6 promotes its inhibition (Lourido et al., 2021). In line with the reduced accumulation of DILP2 in the IPCs of the CerS mutants when fed on HSD, we detected a tendency towards less pronounced downregulation of upd2 (Fig 4L) and less pronounced upregulation of dilp6 (Fig 4M).

In sum, these results lead us to assume that besides the CerS catalytic domain, the CerS Schlank homeodomain might also be a putative target to attenuate the insulin resistance in response to sugar-rich environments.

### CerS NLS2 homeodomain and catalytic domain mutations lead to an altered lipid profile, decreasing sphingolipids in FB upon an HSD

Just as we had done for the KINLS1 and KIE118A CerS homeodomain mutants, we now expressed the NLS2 or H215D variants in FB and examined how these mutations impact the lipid profile when fed on ND or HSD.

First, we assessed the accumulation of neutral lipids in FB lipid droplets by staining the tissue with Nile red. The fat accumulation in terms of lipid droplet area was comparable between the mutants and the wildtype control when fed on HSD (Fig 5A-A’). This is in accordance with the reduced mRNA transcript levels of lipases Vaha (cg8093), cg6277, and lip3 (Fig 5B-D), and the increased TG levels found in the whole larval body and FB (Fig 5E and F). Apart from that, the expression levels of lsd-1 in the FB of these CerS mutants were significantly less upregulated, while the expression of pepck was slightly but not significantly reduced upon HSD (Fig 5G, H). This would explain the low mobilization and high storage of lipids, and the hyperglycemia in the FB caused by an HSD (Lourido et al., 2021; Binh et al., 2019; Zheng et al., 2016).

**Fig 5.**
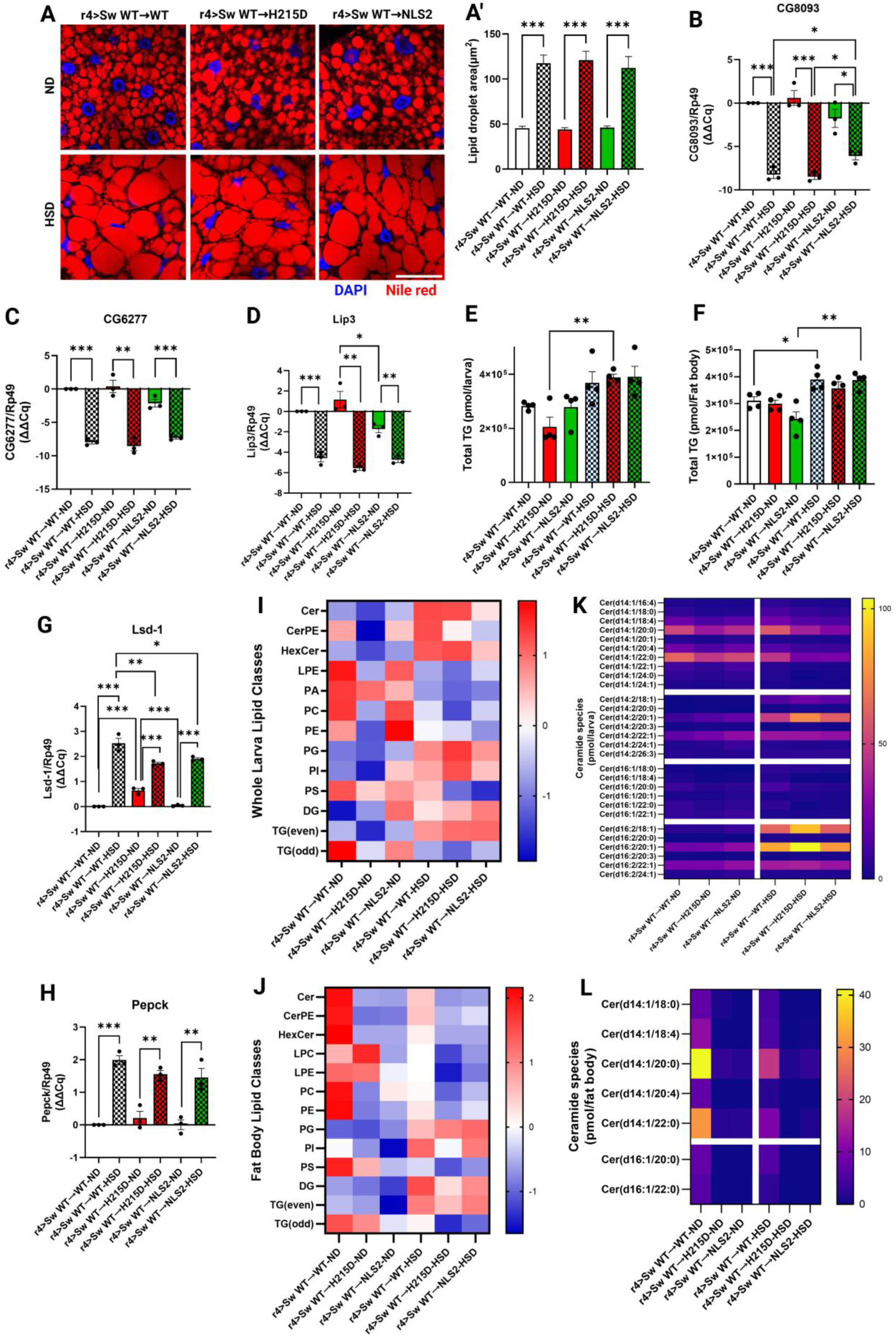
Drosophila CerS Schlank homeodomain and catalytic domain mutations in FB adjust the lipid profile, decreasing sphingolipids in FB upon an HSD. (A) Neutral lipid (Nile Red) staining of FB from larvae bearing CerS mutations in FB, reared on HSD or ND, compared with wild-type control. (A’) Quantification in two-dimensional confocal slices of lipid droplet area in (A). The FB CerS mutants and wildtype L3-instar larvae showed a high increase of lipid droplet area upon HSD compared to the animals reared ND (n≥15). (B-D) Relative expression by RT-qPCR of lipases (B) cg8093, (C) cg6277, and (D) lip3 in the whole larval body of animals bearing CerS mutations in FB compared to the wildtype, showing their significant downregulation in animals reared on HSD (n=3 biological replicates). (E) Total TGs extracted from whole larvae upon ND or HSD and measured by tandem shotgun MS, indicating an upregulation of total TGs when animals are reared HSD (n=4 biological replicates). (F) Total TGs extracted from FB of larvae bearing FB CerS-mutated variants and wild-type animals upon ND or HSD, measured by tandem shotgun MS, indicating an upregulation of total TGs when animals are reared HSD (n=4 biological replicates). (G-H) Relative expression by qPCR of lsd-1 and pepck in FB of larvae fed HSD. It is shown significantly less upregulation of (G) lsd-1 and slightly less but not significant upregulation of (H) pepck in FB from larvae expressing CerS catalytic domain and homeodomain mutations reared on HSD compared to wild-type (n=3 biological replicates). Expression levels are normalized on rp-49, and the ΔΔCq is represented and used in statistical analysis. (I) Heatmap summarizing the representative lipid classes quantified relative to lipid standards in each replicate of one larval body per group and standardized into z-score from whole L3-instar larvae bearing FB CerS-mutated variants and wildtype reared on ND and HSD. Lipids were extracted and measured by tandem shotgun MS (n=4 biological replicates). (J) Heatmap summarizing the representative lipid classes quantified relative to lipid standards in each replicate of one FB per group and standardized into z-score from the whole body of L3-instar larvae bearing FB CerS-mutated variants and wildtype larvae reared on ND and HSD. Lipids were extracted and measured by tandem shotgun MS (n=4 biological replicates). (K) Heatmap representation of ceramide species extracted from whole L3-instar larvae bearing FB CerS-mutated variants and wildtype larvae reared ND and HSD and measured by tandem shotgun MS (n=4 biological replicates). (L) Heatmap of ceramide species extracted from FB of larvae bearing FB CerS-mutated variants and wildtype animals reared on ND or HSD, measured by tandem shotgun MS (n=4 biological replicates). Abbreviations: Ceramide (Cer), Ceramide phosphatidylethanolamine (CerPE), Hexosylceramide (HexCer), Diacylglyceride (DAG), Lyso-phosphatidylcholine (LPC), Lyso-phosphatidylethanolamine (LPE), Phosphatidylcholine (PC), Phosphatidylethanolamine (PE), Phosphatidylglycerol (PG), Phosphatidylserine (PS). Error bars are ± SEM. Unpaired two-tailed t-test, one-way and two-way ANOVA, followed by Tukey’s multiple comparisons tests, were used to derive P-values;* p<0.05, ** p<0.01, *** p<0.001.

Next, we profiled lipids of the whole larval body from controls and animals expressing NLS2 or H215D in the FB, focusing again on SLs and PLs (Fig 5I). Analysing the lipidome, we observed on ND a generally strong reduction of SLs and PLs in H215D mutants as expected when CerS activity is abolished. In contrast, the lipid levels of NLS2 mutants were comparable to the wild-type. On HSD, we detected an increase of SLs (Cer from ∼252,86 pmol/larva on ND to ∼505,02 pmol/larva, and HexCer from ∼124,19 pmol/larva on ND to ∼430,92 pmol/larva in average) in the catalytically inactive H215D mutants and wild-type control (Cer from ∼313,17 pmol/larva on ND to ∼507,65 pmol/larva, and HexCer from ∼206,11 pmol/larva on ND to ∼419,61 pmol/larva in average). Strikingly, this SL increase is much less pronounced in NLS2 animals (Cer from ∼338,08 pmol/larva on ND to ∼407,31 pmol/larva, and HexCer from ∼193,96 pmol/larva on ND to ∼332,70,61 pmol/larva in average). PL levels were quite similar, except for the reduction of PS in the mutants compared to the control (Fig 5I). Furthermore, we detected here, as noted above for KIE118A, KINLS1, and control animals (Fig 3J, L; S4 Fig) a shift of the SL profile towards longer, doubly unsaturated sphingoid bases on HSD. In complete larvae, we found a marked increment in both specific Cer and HexCer species containing sphingoid bases with two double bonds, such as d14:2/18:1, d14:2/20:1, d16:2/18:1, and d16:2/20:1 (Fig 5J; S5E, E’, I, I’ Fig). Conversely, some Cer and HexCer species containing monounsaturated sphingoid bases were decreased upon HSD, such as d14:1/20:0 and d14:1/22:0 (Fig 5K; S5I Fig). Noteworthy, it was recently shown that loss of peroxisomes shifted the SL profile towards longer sphingoid bases and this shift was connected with impaired DILP2 secretion (König et al., 2025).

Now we analysed the lipid profile of NLS2 or H215D variants in the FB. We observed a general lipid decrease with the exception of diacylglycerides (DG), phosphatidylglycerol (PG), and even TGs after a long-term ND and HSD (Fig 5J). Of note, Cer levels were particularly low in the FB of the H215D (from ∼5,13 pmol/fat body on ND to ∼1,14 pmol/fat body on HSD in average) and NLS2 (Cer from ∼3,55 pmol/fat body on ND to ∼4,21 pmol/fat body on HSD in average) variants, in comparison to wildtype (Cer from ∼111,9 pmol/fat body on ND to ∼47,07 pmol/fat body on HSD in average) (Fig 5L). This result is expected for the catalytical inactive H215D mutants. For the NLS2 mutation the extent of reduction of Cer species in the FB was unexpected since no effect on Cer levels was found here (Fig 5 G) or when CerS activity was analysed in complete larvae in an earlier study (Sociale et al., 2018). However, reduced Cer content in the FB was already observed for the KIE118A and KINLS1 mutants under an ND regime and in mammals for CerS2 E115A variant (Nicholson et al., 2021). This suggests that mutations in the homeodomain of CerS proteins may have a general impact on Cer production.

Nonetheless, although Cer levels in these FB-expressed CerS mutants were very low, we found in wild-type animals a change of Cer species similar to what we observed in the CerS KIE118A and KINLS1 homeodomain mutants and their wild-type control. The species Cer (d14:1/20:0) and Cer (d14:1/22:0), with a single double bond sphingoid base were strongly decreased in animals fed on HSD (Fig 5L, S5D Fig). This decrease could very well be connected with a shift of the SL profile towards longer, doubly unsaturated sphingoid bases on HSD, which we cannot detect here in FB due to the generally very low ceramide levels. Nonetheless, ceramides are greatly reduced, suggesting, in line with data from earlier studies, that lowered ceramide level resolves insulin resistance. In sum, these data suggest that reducing Cer levels via mutation of the catalytic domain or the specific mutation in the homeodomain leads to amelioration of insulin resistance, making the homeodomain an attractive therapeutic target.

## Discussion

Several studies have been conducted to address the pathogenesis of diabetes mellitus, including the study of lipid trigger(s) such as ceramides. To date, inhibitors of the SL biosynthesis pathway, specific CerS inhibitors and the multi-target designed ligands (MTDLs) have been used to tackle the C16- and C18-ceramides associated with insulin resistance among other diseases (Alizadeh et al., 2023; Schiffmann et al., 2012). However, those studies are mainly focused on the CerS enzymatic activity, neglecting the function of its homeodomain. Although recent studies have shown that the CerS homeodomain possesses transcriptional regulation properties (Sociale et al., 2018; Yuan et al., 2024), its function under a hypercaloric diet regimen remains poorly understood. This study demonstrates that the Drosophila CerS homeodomain, beyond its catalytic lag1p motif, is crucial for regulating lipid content and, notably, insulin sensitivity during caloric overload, such as on HSD. Through a combination of genetic tools, immunostaining and lipidomics, we found that targeting the catalytic domain, as well as various sites in the homeodomain, including the two nuclear localization sites (NLS1 and NLS2) and an E118A substitution resembling a human SNP (E115A), leads to distinct consequences. While KINLS1 and KIE118A mutations predispose, to some extent, to insulin resistance when expressed ubiquitously, the conditional cis-alleles switch NLS2 mutation instead mitigates insulin sensitivity loss when expressed in the FB, showcasing a tissue-specific effect. These results build on previous findings in mammals, where the missense mutation E115A negatively affected CerS2 catalytic activity and enhanced the glucose intolerance and hepatic steatosis when placed on the obesogenic diet (a high-fat diet; HFD) in mice, although it was insufficient to affect the serum SL profile in humans (Heinitz et al., 2024; Nicholson et al., 2021). Moreover, this data builds on our prior work concerning the CerS homeodomain, which presents the NLS2 involved in DNA-binding and transcriptional regulation (Sociale et al., 2018). This study, therefore, demonstrates that each mutation within the CerS homeodomain has particular consequences on sphingolipid biosynthesis, insulin secretion and sensitivity.

### Drosophila CerS *human SNP* and NLS1 mutation in the homeodomain predispose to loss of insulin sensitivity and a shift in the lipidome on HSD

Firstly, the production and secretion of insulin/DILP is crucial in metabolic homeostasis across vertebrate and invertebrate species. In Drosophila, the IPCs are the main source of DILP2, 3 and 5, although DILP1 is also produced transiently during the pupa stage until a few days of adult life (Liu et al., 2016; Semaniuk et al., 2021). By ubiquitously expressing the E118A and NLS1 knock-in in an HSD T2D Drosophila model, we demonstrated that there is a disruption in the release of DILP2 from the IPCs, leading to the accumulation of DILP2 and susceptibility to insulin sensitivity loss. The level of these DILPs is influenced by various external factors, including nutrition, circadian rhythm, temperature, drugs, or genetic manipulation. Besides, their biosynthesis and secretion depend on internal factors, such as peptides produced in the periphery (e.g. the production of upd2 and dilp6 in the FB), that might modulate DILP production and/or release by targeting IPCs activity (Nässel et al., 2013; Nässel & Broeck, 2016; Semaniuk et al., 2021). Previous studies and this research have shown that under chronic HSD consumption, the DILP peptides are accumulated in the IPCs of Drosophila larvae, resembling the alteration of the synthesis and release of insulin in a mouse model (de María Márquez Álvarez et al., 2023; Khan et al., 2025; Márquez Álvarez et al., 2024; P et al., 2020). Although the brain mRNA levels of dilps were increased after an HSD regimen, no significant difference was observed between the CerS mutants and wildtype control. However, the accumulation of DILP2 in the IPCs was notably increased in the KIE118A mutant. This suggests that the failure in DILP2 secretion is particularly enhanced when the Drosophila CerS homeodomain is bearing a mimicked human SNP. Furthermore, we confirmed the presence of insulin resistance by assessing the expression of the insulin resistance marker nlaz in the FB, which was upregulated after HSD consumption in both CerS mutants and wildtype control. What is more, the localization of FoxO in the FB was certainly nuclear, reaffirming the insulin resistance and the loss of insulin sensitivity in CerS mutants and wildtype under this caloric overload condition. Nonetheless, the nuclear localization of the KIE118A mutant was higher, implying that KIE118A has an effect in both the IPCs and the FB to disturb the secretion of DILP and enhance the shuttling of FoxO to the nucleus, respectively (Dobson et al., 2017; Lee & Dong, 2017;Zang et al., 2024; Zhao et al., 2021). Two peptides, UPD2 and DILP6, expressed in the FB and already known to be targeting the IPCs in an organ-to-organ manner were tested (Nässel et al., 2013). Although the expressions of upd2 and dilp6 in the FB were not significantly different between the CerS mutations and wildtype upon an HSD, the downregulation of upd2 and upregulation of dilp6 are consistent with the accumulation of DILP2 in the IPCs (Bai et al., 2012; Ingaramo et al., 2020; Semaniuk et al., 2021). While the KINLS1 mutant showed a mild effect on the insulin sensitivity, the CerS KIE118A mutant showed a strong effect on the loss of insulin sensitivity. Therefore, it is conceivable to hypothesize that the KIE118A mutant, apart from influencing the shuttling of FoxO, might be disturbing other crucial unknown factors released from the FB to the systemic circulation, targeting the IPCs for the release/retention of DILP in an interorgan communication manner.

On the other hand, as shown by others (Musselman et al., 2011; P et al., 2020) and by this study, the chronic consumption of HSD leads to fat accumulation in the FB. In addition, the lipolytic regulation is mainly carried out on the surface of the lipid droplet by various lipases when the release of fatty acids from TGs is required (Nirala et al., 2013; Singh et al., 2024; Zheng et al., 2016). Among the downregulated lipases assessed in this study, the secretory lipase CG8093 called “Vaha”, showed up as a recently shown factor that plays an important role in being released to the systemic circulation and targeting the IPCs, stimulating the secretion of DILPs (Singh et al., 2024). Although no significant difference was found in the expression of genes involved in lipolysis and lipogenesis, the low expression of cg8093 when animals are under HSD regimen indicates less DILP release. This also might support the assumption that a complex network of several secretory peptides regulates the secretion of DILPs and IPCs activity.

Analysing the lipidome, we observed a decrease in total ceramide in the FB, but not in the whole larval body of CerS KIE118A and KINLS1 mutants upon an ND. Conversely, upon an HSD, the total ceramide was slightly increased in the whole larval body, while it decreased in FB; nonetheless, the CerS mutants presented slightly more total ceramide in FB compared to the wildtype control. This suggests that the KIE118A and KINLS1 mutations have only a modest impact on enzyme activity similar to what was found for the human SNP variant in liver homogenate (Nicholson et al., 2021).

Aside from that, we notably detected changes in specific ceramide species in both the FB and the whole larval body when the animals were reared on HSD. Whereas the CerS2 E115A mutant mice demonstrated increased synthesis of long acyl chain C16-ceramide under HFD consumption (Nicholson et al., 2021), we noted that after a long-term HSD consumption in Drosophila, specific Cer and HexCer species containing sphingoid bases with two double bonds were elevated. Additionally, other ceramide species containing monounsaturated sphingoid bases were decreased. Recent studies in mammals have revealed that apart from the dihydroceramide desaturase 1 (DEGS1), which introduces a ΔC4,5 trans-double bond (Δ4E) into the sphinganine backbone, the fatty acid desaturase 3 (FADS3) may introduce a kinked second cis double bond (Δ14Z) in the sphingoid base, likely influencing membrane biophysical properties such as fluidity and lipid-protein interactions, thereby affecting the cell signaling (Hornemann, 2025; Jojima et al., 2020; Jojima & Kihara, 2023; Karsai et al., 2020). This finding suggests that the additional double bond in the sphingoid base might be a contributing factor to the increased risk of insulin resistance caused by a hypercaloric diet (Chaurasia et al., 2019; Hornemann, 2025). Thus, it is highly likely that the production and distribution of ceramide containing mono- or polyunsaturated sphingoid bases might depend on the caloric load of the diet. The balance in the unsaturation number of ceramide sphingoid bases could be critical in the composition and maintenance of the lipid membrane.

### Targeting Drosophila CerS NLS2 homeodomain and catalytic lag1p motif function in FB alleviates the insulin resistance upon an HSD

Multiple studies including ours show that deletion of CerS or inhibition of CerS activity attenuates the metabolic disorder caused by caloric overload (Hammerschmidt et al., 2023; Khan et al., 2025; Raichur et al., 2014, 2019; Turpin et al., 2014). However, most of these studies focus solely on enzymatic activity. We recently found that the CerS in Drosophila regulates lipid metabolism not only through its enzymatic activity, but also by its homeodomain as transcriptional regulator (Sociale et al., 2018; Voelzmann et al., 2016; Yuan et al., 2024). Ubiquitous expression of the catalytic inactive H215D variant or of the NLS2 variant, which leads to loss of DNA binding result in early lethality. To examine of the H215D or of the NLS2 variant in insulin resistance we have generated new lines switching from a wild-type CerS to wild-type HA-tagged CerS (control) or to either H215D- or the HA-tagged NLS2-variant, which we expressed in FB. The Drosophila FB is a multifunctional tissue that responds to various stimuli and plays a prominent role in regulating the insulin secretion from the neurosecretory IPCs. In addition to the characteristic effects of a chronic HSD consumption, such as hyperglycemia, reduced larval size and weight, and a delayed developmental rate, we observed notably less nuclear localization of FoxO in the FB of both CerS mutants. Surprisingly, this effect was more pronounced in the NLS2 homeodomain mutants compared to the catalytic domain mutants. Even when the ratio of nuclear/cytoplasmic was not completely recovered, as in ND conditions, the inhibition of the CerS enzymatic activity and, interestingly, the transcriptional regulation seemingly improves insulin sensitivity. Also, the lower upregulated expression of nlaz in the CerS mutants could explain the mild alleviation of insulin resistance in the FB.

As mentioned in a previous section, sufficient evidence has thus far linked the function of the FB and IPCs to the production and release of insulin, which accumulates in IPCs following a chronic HSD consumption (Semaniuk et al., 2021). Despite the upregulated mRNA levels of the dilps in the IPCs after an HSD, the CerS mutants presented slightly lower expression of dilps. Furthermore, we unexpectedly observed an enhanced secretion of DILP2 from the IPCs in both mutated variants of CerS. Given that the FB acts as a sensor of nutritional state, and its signaling factors may influence the expression and secretion of one or more DILPs, we detected in the CerS mutants a couple of signaling factors that target the IPCs and regulate the release of DILPs (Bai et al., 2012; Ingaramo et al., 2020; Rajan & Perrimon, 2012; Zheng et al., 2016). We saw less downregulated expression of upd2 and less upregulated expression of dilp6 as compared to the wildtype control. Although the expression of these factors was not completely restored to control levels as in animals on ND, it is likely that a variety of other factors, not yet identified, might interact with CerS. Thus, we propose that targeting not only the catalytic domain but also the NLS2 motif in the homeodomain of CerS might play a role in a complex intracellular regulation of signaling factors produced and secreted from the FB to the systemic circulation. However, further investigation is required to understand the mechanisms and implications of CerS in organ-to-organ communication.

On the other side, expression profiling studies have suggested that lipogenesis, lipid storage, and beta-oxidation are increased after chronic HSD consumption (Musselman et al., 2011, 2013). Although no significant differences were found in the high lipid accumulation in lipid droplets of FB among the animal groups when reared on HSD, we detected a significantly lower upregulation of lsd-1, as well as no significant but slightly lower upregulation of pepck in the CerS mutants. Even though the expression levels of these factors did not completely restore to those detected in animals on ND, they were mildly improved after an HSD in CerS mutants. Of note, this effect is accompanied by a strong impact on the FB lipidome, reducing total lipid content and likely helping facilitate the mobilization of lipids.

It is widely accepted that the N-acyl chain length of ceramides may determine the effect on the metabolic rate, leading to lipotoxicity and triggering metabolic disorders upon their imbalance. Several studies have unveiled that the increment of long-chain C16- and C18-ceramide may trigger glucose intolerance and insulin resistance upon an obesogenic diet in mammals (Bandet et al., 2019; Chitkara & Atilla-Gokcumen, 2025). It was also suggested that very long-chain C22- and C24-ceramide might be playing a protective role in this regard (Bandet et al., 2019; Turpin et al., 2014; Turpin-Nolan et al., 2019). However, ultra-long-chain C-40:1 and C-42:1 ceramide, found in murine and human skeletal muscle, might be associated with lipotoxicity and metabolic disease (McNally et al., 2022). Conversely, our lipidomic data from the mutated catalytic motif and NLS2 homeodomain reveal that whereas specific ceramide and HexCer species containing sphingoid bases with two double bonds were elevated upon a hypercaloric diet, other specific ceramide sphingoid bases with a single double bond were decreased in the whole larval body but not in the FB. Intriguingly, this tendency was also observed in the lipidomic analysis shown in a previous section of the CerS homeodomain mutated variants, KIE118A and KINLS1, in the whole larval body and the FB. The effect of consuming HSD resembles a recent study, where the loss of peroxisomes caused by mutations in the peroxisome assembly factor Peroxin Pex19 shifts the profile of SL towards longer sphingoid bases with doubly unsaturated chains. Additionally, since the peroxisomes interact with the Golgi apparatus in IPCs, when Pex3, Pex16 or Pex19 are abolished, the secretion of DILP2 from IPCs is impaired (König et al., 2025). In a previous study, it was demonstrated that Pex19 was associated with Drosophila CerS Schlank. By lacking peroxisomes, the fatty acids cannot be oxidised, leading to a reduced production of medium-chain fatty acids (MCFAs), which play a crucial role in triggering the shuttling of CerS Schlank homeodomain to the nucleus, thereby repressing lip3. When CerS Schlank is not shuttled to the nucleus, lip3 is derepressed, promoting the production of FFA, which in turn activates HNF4 signalling and a pathological cascade (Bülow et al., 2018; Sellin et al., 2018). Since Golgi membranes are rich in ceramides (König et al., 2025), it is conceivable hypothesize that Drosophila CerS might modulate the packing of DILPs into dense-core vesicles at the trans-Golgi network, not only by its enzymatic function but also by its homeodomain.

Even when a previous study showed that the NLS2 mutation did not affect the catalytic activity when expressed ubiquitously in the complete larva (Sociale et al., 2018; Voelzmann et al., 2016), the ceramide species found in both the FB-expressed CerS homeodomain NLS2 and catalytic domain H215D mutated variants were drastically reduced. Furthermore, not only ceramides containing monounsaturated sphingoid bases were reduced, but also ceramides containing sphingoid bases with a second double bond were not detected. Since Drosophila CerS is expressed ubiquitously, it is unclear whether the effect of the NLS2 mutation can be replicated in other tissues beyond the FB.

The reduction in the nuclear localization of FoxO and improvement in the release of DILP2 from the IPCs in the FB-expressed CerS mutants after an HSD regimen could be explained by their low production of SL species in the FB. Thus, this outcome suggests that reducing the shift of SL towards longer, doubly unsaturated sphingoid bases in the FB on a long-term HSD consumption, might diminish the overall content of this type of sphingoid bases, and as a consequence ameliorating the insulin resistance. In addition, two double-bond sphingoid bases might interact with intracellular secretory and non-secretory factors, which could be involved in lipotoxicity, impacting on insulin sensitivity (Chaurasia et al., 2019; Hornemann, 2025). As mentioned above, the FADS3 may introduce a second double bond (Δ14Z) in the sphingoid base. Although the impact of a second double bond in the sphingoid base on individuals under a hypercaloric diet regimen is not well understood, this finding may encourage further studies to identify a new biomarker for consideration in insulin-related metabolic disorders.

In summary, this study provides insights into the role of CerS in insulin-associated metabolic disorders, emphasizing on the homeodomain rather than the catalytic domain. Importantly, our findings demonstrate that the nuclear localization sites NLS1 and NLS2, as well as the E118A substitution, which mimics the human SNP E115A variant, result in individual repercussions on the insulin-related metabolic processes under a high-sugar diet regime. While the ubiquitously expressed CerS E118A and NLS1 mutations appear to increase the risk of insulin resistance, in contrast, the FB-expressed NLS2 mutation mildly alleviates the insulin resistance. Due to the improving effect of the FB-expressed catalytic domain and homeodomain mutants on high sugar-mediated insulin resistance, we encourage further studies of the CerS mutated variants expressed in other Drosophila tissues to understand the behaviour of both its enzymatic production and its homeodomain in transcriptional regulation. Likewise, due to the limited information about the homeodomain function in mammalian CerS under an obesogenic diet regime, future work in a mouse model should include the study of the nuclear localization site NLS2 and its impact on enzymatic activity.

## Materials and Methods

### Schlank Alleles

Schlank mutant alleles were generated as we have done earlier (Sociale et al., 2018). Briefly, a founder schlank knockout (KO) line, w-schlank^KO^ with an attP site and a single loxP site at the schlank endogenous gene locus was generated by homologous recombination as described (Huang et al., 2008). Target constructs containing the deleted genomic DNA (gDNA) of schlank with a point mutation in NLS1 (pGE-attB [KINLS1]) and E118A (*pGE-attB* [KIE118A]) lying within the first and third helix of the homeodomain (Fig 1A), respectively, were reintegrated by ΦC31-mediated integration into the attP site of the *w-*, *schlank^KO^*founder line. Similarly (pGE-attB [KIH215D]) was generated.

Just like we had done before (Sociale et al., 2018) for the founder knock-out lines (KO lines) we tested for complementing or non-complementing the schlankG0349 mutant allele (See Complementation Tests) which was previously generated (Bauer et al., 2009).

To enable cis-allele switching, target constructs containing the wildtype variant of deleted gDNA of schlank together with a second wild-type variant of Schlank gDNA (RIV FRT.KIWT.pA.FRT.KIWT.HA) or gDNA containing either a point mutation within the NLS2 site (RIV FRT.KIWT.pA.FRT.KINLS2.HA) or at amino acid position 215 within the catalytic lag motif (RIV FRT.KIWT.pA.FRT.KIH215D.HA) were reintegrated by ΦC31-mediated integration into the attP site of the *w^-^*, *schlank^KO^* founder line. Excision of the FRT cassette makes the locus stop expressing wild-type schlank and start expressing HA-tagged Schlank variants instead. In the Fig 4 and Fig 5 named as Sw WT→WT, Sw WT→NLS2, and Sw WT→H215D respectively.

All ΦC31-mediated integration experiments into our *w-schlank^KO^*founder line were done by BestGene Inc. (Chino Hills, CA, USA).

### Complementation Tests for The Engineered *schlank* Alleles

*schlankKO/FM7a, P{Dfd-GMR-nvYFP}1* and *schlankKIH215D/FM7a, P{Dfd-GMR-nvYFP}1 n* were generated by crossing these lines with *w, schlankG0349/Dp(1;Y)dx+1/C(1)M5* males. Transheterozygous *schlank* female progeny are larval lethal with both alleles. *schlankG0349/FM7a, P{Dfd-GMR-nvYFP}1 were crossed with KINLS1/Y or KIE118A/Y males.* Transheterozygous *schlank* female progeny are non-lethal with both alleles.

### Fly Strains

pP{UAS-schlank-CTHA}, WT-HA and pP{UAST-Schlank NLS1 HA}, NLS1-HA (Voelzmann et al., 2016); pP{UAST-Schlank H215D-HA} (Sociale et al., 2018); r4-GAL4 (Bloomington #33832)

### Plasmids and Cloning

The deleted 2,4 kb gDNA of schlank containing exons 2 to 6 and the 3’UTR region was amplified via PCR and cloned into pSC-A-amp/kan plasmid (Topo TA Cloning Kit, StrataClone) using XhoI und NheI. Mutations within target constructs KINLS1 (NLS1 RPKK to RPMM), E118A (E118 to 118A), and H215D were introduced via mutagenesis PCR. The gDNA fragments were cut out and cloned into the pGE-attB integration vector bearing a *w+* marker using XhoI und NheI restriction sites.

Wild-type rescuing Schlank-gDNA was cloned into the multiple cloning site (MCS) of the reintegration vector (RIV FRT.MCS.pA.FRT.MCS3 vector) (Baena-Lopez et al., 2013), which is flanked by two FRTs to render it excisable by FLP using XhoI und NheI. Into the MCS3 site, another gDNA downstream of the one flanked by FRTs was inserted. The gDNAs inserted into MCS 3 were: wild-type Schlank-gDNA-HA, HA-tagged Schlank-gDNAs carrying the NLS2 or the H215D mutation using PacI and AgeI finally resulting in RIV FRT.KIWT.pA.FRT.KIWT.HA, RIV FRT.KIWT.pA.FRT.KINLS2.HA, and RIV FRT.KIWT.pA.FRT.KIH215D.HA. This configuration can be used for cis-allele switching, whereby the wild-type Schlank gDNA can be replaced by a wild-type or mutant allele at specific times and places.

NLS1-HA and E118A-HA double-stranded, sequence verified genomic block (gBlock) containing the entire Schlank cDNA with a mutation within the NLS1 motif or with a mutation in E118 were generated by Integrated DNA Technologies, Inc. (IDT). The fragments were inserted into pP{UAST} using EcoRI and XhoI.

### Fly Diet Preparation

Stocks were maintained on a standard cornmeal-yeast-agar medium. For experiments, a nutritionally balanced diet (Control diet, 0.15M sucrose) was used as a control to assess the effect of high-sugar diets (HSD, 1M sucrose) on fly metabolism. The total nutritional values are 702.6 kcal for ND and 1866.6 kcal for HSD. Food cooking was performed as follows: Agar, brewer’s yeast, Peptone, yeast auto lysate, sugar, MgSO4 x 6H2O, and CaCl2 x 2H2O were added to a beaker and mixed with a small amount of water (bidest.). The mixture was heated in a microwave until the ingredients were fully diluted, after which more water was added till the wanted volume was achieved. The mixture was heated again until boiling. The mixture was then cooled to 50 °C, after which Nipagin (10% in ethanol) and propionic acid (10% in ethanol) were added, while continually stirring. The media were then filled into small culture plates, where they would harden, and were stored at 4°C until further use. The diets were prepared according to protocols previously described (Musselman et al., 2011).

### Selection of Larvae for Analysis, Pupation and Survival Assay

For the experiments, only male larvae were taken. Females of homozygous stocks were crossed to *FM7a, P{Dfd-GMR-nvYFP}* carrying males. Eggs were collected for 4 h on apple juice agar plates, and L1 instar larvae were sorted for the absence of the dfd-nvYFP marker and transferred to a new plate. CerS Schlank mutants, as well as *w^1118^*, KIWT or Sw WT→WT (used as control) larvae were staged as developmentally comparable with 96 h when their anterior spiracles had not yet protruded from the cuticle and when posterior spiracles turned to dark orange.

For the pupation and survival assays, 25 larvae were counted and monitored throughout the larval and pupal stages. In particular, KINLS2-mutated pupae were transferred to a petri dish containing a filter paper soaked in PBS to avoid decomposing during the slow development. The experiments were performed in triplicate as a minimum.

For experiments with a normal diet (ND) and high sugar diet (HSD), embryos were collected on apple juice agar plates, and 24 h AEL, the early first instar larvae were transferred to a ND or HSD until 72 h and 120 h AEL, respectively, at 29 °C. For comparative purposes, the third instar larvae (L3) were selected as morphological criteria to ensure that WT and mutant animals were at the same developmental stage.

### Larval body length and wet weight measurement

Larvae were collected, rinsed thoroughly with 1X PBS to remove any remaining food and dried on tissue paper. For the length measurement, larvae were transferred to a 1.5 ml tube containing 1x PBS, heated to 65°C for approximately 3 minutes, and measured along the anterior-posterior axis on a glass slide. For the wet weighing, larvae were collected in groups of 10 in a 1.5 ml tube using an analytical balance.

### Metabolic Labelling of Lipids of Larvae

TG, fatty acid, and CerS activity were determined as described by measuring de novo TG, fatty acid, and ceramide generation after 18 hr labelling of larvae with [1-14C]-acetic acid. For metabolic labelling, larvae were fed with inactivated yeast paste containing [1-14C]-acetic acid (61 Ci/mol) as a C1-precursor for lipids. The procedure was carried on as previously described (Bauer et al., 2009).

### Lipidomics

Lipid extraction: Lipids were extracted from individual larvae. Each larva was homogenized in 100µL water (LC-MS grade). To every homogenate, 1000µL Extraction mix (5/1 MeOH/CHCl3 containing internal standards. PE[31:1] 420pmol; PC[31:1] 792pmol; PS[31:1] 197pmol; PI[34:0] 169pmol; PA[31:1] 112pmol; PG[28:0] 103pmol; CL[56:0] 57pmol; LPA[17:0 79pmol; LPC[17:1] 70pmol; LPE[17:0] 76pmol; Cer[17:0] 64pmol; SM[17:0] 198pmol; GlcCer[12:0] 110pmol; GM3[18:0-D3] 29pmol; TG[50:1-D4] 2351pmol; CE[17:1] 223pmol; DG[31:1] 128pmol; MG[17:1] 207pmol; Chol[D6] 1448pmol; Car[15:0] 91pmol; Sph[d18:0-D7] 64pmol) were added and tubes were sonicated for 30 minutes in a bath sonicator. After 2 minutes of centrifugation at 20000g, supernatant was transferred to a fresh Eppendorf tube. 200 µL chloroform as well as 600µL 1% acetic acid were added to each tube, tubes were shaken manually for 10 seconds followed by centrifugation for 5 minutes at 20000g to separate phases. The lower phase was transferred to a fresh tube and tubes were dried in a speed-vac for 15 minutes at 45°C. Dried samples were redissolved by adding 1mL of spray buffer (8/5/1 isopropanol/methanol/H2O (all LC-MS grade) + 10mM ammonium acetate + 0.1% acetic acid (LC-MS grade)) to each sample and sonicating for 5 minutes in a bath sonicator. Mass spectra were recorded on a Thermo Q-Exactive Plus spectrometer equipped with a standard heated electrospray ionization (HESI) II ion source for shotgun lipidomics. Samples were sprayed at a flow rate of 10 µl min−1 in spray buffer. MS1 spectra (res. 280000) were recorded as segmented scans with windows of m/z 100 between m/z 240 and 1,200 followed by MS2 acquisition (res. 70000) from m/z 244.3364 to 426 1,199.9994 at m/z 1.0006 intervals. Raw files were converted into mzml files and analysed using 427 the LipidXplorer (version 1.2.8) ((Herzog et al., 2011)). Internal standard intensities were used to calculate 428 absolute amounts, and all identified lipids were normalized to the total lipid amount.

### Cell Culture and Luciferase Assays

Assays were done as described earlier (Sociale et al., 2018). Drosophila Schneider cells (S2) were grown in Schneider’s medium supplemented with 10% heat-inactivated fetal bovine serum and 1% penicillin/ streptomycin (Sigma-Aldrich, USA). Transfection was done 12 hr after plating S2 cells (Effectene; QIAGEN) according to the manufacturer’s instruction. Starvation and fatty acid treatment were done in Serum Free Optimem with or without BSA-fatty acids for the indicated times. Luciferase assays were performed 48 hr after transfection of the corresponding plasmids with the Dual-Luciferase Reporter Assay System (Promega) according to the manufacturer’s protocol. Activity was determined using a MicroLumatPlus LB 96V luminometer (Berthold Technology). Each experiment was repeated three times and is the average of the reading of three different transfected wells. Verification of efficacy and specificity of the lip3 reporter was tested as described in (Sociale et al., 2018).

### RNA Extraction and RT-qPCR

The total RNA was extracted from the whole larvae, brain or FB using TRIzol Reagent (Invitrogen™, Carlsbad, CA) following the manufacturer’s instructions. RNA concentration was measured with the Thermo Scientific™ NanoDrop™ 2000. The cDNA was synthesised by the LunaScript® RT Super Mix Kit from New England Biolabs®. The RT-qPCR was performed in a reaction mix containing cDNA template, primers, and the Luna® Universal qPCR Master Mix from New England Biolabs® or iQ SYBR Green Supermix (Bio-Rad). The experiments were performed in a CFX Connect Real-Time PCR Detection System (Bio-Rad) or an iQ5 RT-PCR. Ribosomal protein L32 (rp49) was used as a reference gene. The experiment was performed using three biological replicates. Primers are described in the Supporting Information (S2 Table).

### ChIP

ChIP was performed according to (Andresini et al., 2016) and as described in detail earlier (Sociale et al., 2018). Briefly, S2 cells were crosslinked with formaldehyde. After sonication of nuclei to yield DNA fragments of about 500 bp immunoprecipitation was performed with the indicated antibodies. Upon reversion of the crosslinking, DNA was extracted with phenol/chloroform, and 1 ml of the immunoprecipitated DNA was used for qRT-PCR analysis. Expanded protocol is provided in the Supporting Information.

### Immunohistochemistry

Larvae were inverse-prepped in ice-cold PBS. The brains and FB were isolated and fixed for 30 minutes in 4% methanol-free formaldehyde at room temperature. After several washes in Phosphate-buffered saline with 0.1-0.3% Triton X 100 (PBT) and incubation for 30 minutes in blocking solution (5% donkey serum in PBT), primary antibodies (anti-DILP2 1:400 in larval brains (Delanoue et al., 2016) and anti-FoxO 1:500 in FB) (P et al., 2020) were incubated overnight at 4°C. Samples were then washed and incubated in a blocking solution as previously described. Secondary antibodies were added for 1.5 h at room temperature, followed by several washing steps. The tissues were mounted in Fluoromount G with DAPI.

### Nile Red Staining

For lipid droplet staining, the FB from L3 larvae was dissected in 1× PBS and fixed in 4% methanol-free formaldehyde in 1× PBS for 20 min at room temperature. The tissues were then rinsed twice with 1× PBS, incubated for 30 min in 0,00002% Nile red (Sigma) diluted in 0.1% PBT, and washed thoroughly with PBS. The tissues were mounted in Fluoromount G with DAPI.

### Image Acquisition and Processing

To quantify DILP2 levels, confocal Z series of the IPCs were obtained using identical laser power and scan settings. Then, the total fluorescent intensity across the IPCs was measured by generating the maximum-intensity projections of the Z stacks. Images were taken with a Zeiss LSM710 confocal microscope at 63x magnification. The FoxO staining and lipid droplet size quantification were analysed with ImageJ (National Institute of Health, USA Schneider, Rasband, Eliceiri 2012). Where needed, brightness or contrast was adjusted for the whole image in Adobe Photoshop. For comparative stainings, images were not post-processed. Image panels were assembled in Adobe Illustrator software or created with BioRender.com.

### Hemolymph glucose assay

Larvae were collected and rinsed, then hemolymph was collected and assayed as described by (Musselman et al., 2011). Briefly, fine forceps are used to wound and bleed 8–10 larvae, which were combined and centrifuged in an inner 0.5ml tube to isolate 1–2 microliters of hemolymph in an outer 1.5ml tube. Then, 1µl of hemolymph was added to frozen 99µl Infinity Glucose Hexokinase assay reagent (ThermoFisher TR15321) in a 96-well plate and incubated for 15 minutes at 37°C. Absorbance was quantified at 340 nm against a glucose standard curve.

### Western blotting

The whole larval tissue or FB was harvested, homogenized, and boiled in 5x Laemmli buffer, RIPA buffer, and protease inhibitors. After the SDS-PAGE, the membrane was blocked and probed in TBS-T + 5% milk and incubated with primary antibodies overnight at 4°C. Primary Antibodies anti-HA tag (high affinity IgG1, Roche, 1:400), and the secondary antibody Donkey anti-rat Ig-HRP conjugated (Jackson Immunoresearch, 1:10000) were used. The membrane was incubated with Pierce ECL blotting substrate (ThermoFisher Scientific) for a short time and the chemiluminescence was detected with BioRad ChemiDoc MP Imaging System.

### Statistical analysis

All data are presented as means ± standard error of the mean (SEM). We used a two-tailed t-test, one-way and two-way ANOVA followed by Tukey’s post-hoc test using GraphPad Prism software version 9.0.2 (GraphPad Software Inc.) with at least three independent biological replicates (n). A p-value lower than 0.05 was considered statistically significant with *p ≤ 0.05, **p ≤ 0.01 and ***p ≤ 0.001.

### Visualization

The illustrations were developed in Adobe Illustrator or created in https://BioRender.com

## Acknowledgments

We thank Pierre Leopold for kindly sharing the anti-DILP2 and anti-FoxO antibodies. We thank Denis Tsverkun, Sophie Ebert, and Robin Anthonipillai for phenotypical analysis on HSD. We thank Thanh-Phuong Nguyen, Ikram Arahouan for their help with cloning allele switch lines.This research was supported by the National Scholarship and Educational Credit Programme of the Peruvian government (PRONABEC) and by the German Academic Exchange Service (DAAD 57403664). Funding to MHB by the German Research Foundation (DFG, project numbers 417982926 and 535112684), from the University of Bonn (TRA Matter and TRA Life & Health INNOVATION grant) and from the Medical Faculty of the University of Düsseldorf (Integration grant).

## Supporting Information

### Supplemental Tables

**S1 Table.** Resources used in this research.

**S2 Table.** Cloning and fly line generation.

**S3 Table.** List of primers used for qRT-PCR.

**S4 Table.** Primers used for qRT-PCR ChiP.

### Supplemental Figure Legend

**S1 Fig**. Genomic engineering of *schlank mutant alleles*.

**S2 Fig.** Phenotypical analysis of *KO and KIN* mutant lines.

**S3 Fig.** Genomic engineering of *schlank alleles for cis-allele switching*.

**S4 Fig.** CerS E118A and NLS1 homeodomain mutants alter lipid profile, predisposing to insulin disturbance.

**S5 Fig.** FB-expressed CerS NLS2 and H215D mutants modify lipid profile, decreasing sphingolipids.

## Supplemental Tables

**S1 Table.** Resources used in this research.

| REAGENT or RESOURCE | SOURCE | IDENTIFIER |
| --- | --- | --- |
| **Antibodies** |  |  |
| Rat anti-Dilp2 | (Géminard et al., 2009) | N/A |
| Rabbit anti-FoxO | Pierre Leopold-Institut Curie | N/A |
| Goat anti-rabbit Ig-HRP | Jackson ImmunoResearch | 111-035-144 |
| Donkey anti-rabbit Cy3 | Dianova | SKU: 711-166-152 |
| Donkey anti-rabbit Alexa Fluor 488 | Dianova | SKU: 711-545-152 |
| **Chemicals, recombinant proteins** |  |  |
| Infinity Glucose Hexokinase assay reagent | Thermo Fisher Scientific | TR15321 |
| Glucose standard solution | Merck | G6918 |
| Nile red | Merck | N3013 |
| Fluoromount G with DAPI | Invitrogen | 00-4959 |
| **Experimental models:**<br>**Organisms/Strains** |  |  |
| *schlank*<sup>KO</sup> | This Paper | N/A |
| *schlank*<sup>KIWT</sup> | This Paper | N/A |
| *schlank*<sup>KINLS2</sup> | This Paper | N/A |
| *schlank*<sup>KINLS1</sup> | This Paper | N/A |
| *schlank*<sup>KIE118A</sup> | This Paper | N/A |
| *Schlank*<sup>FRT.KIWT.FRTKI.WTHA</sup> | This Paper | N/A |
| *Schlank*<sup>FRT.KIWT.FRTKI.HDHA</sup> | This Paper | N/A |
| *Schlank*<sup>FRT.KIWT.FRTKI.NLS2HA</sup> | This Paper | N/A |
| P{r4-GAL4}3 | FlyBase | Bloomington Nr.33832 |

**S2 Table.** Cloning and Allele Switch fly line generation. The fly lines *Schlank Sw WT*→*WTHA, Schlank Sw WT*→*H215DHA and Schlank Sw WT*→*NLS2HA* were generated by homologous recombination and φ-mediated integration (BestGene Inc., Chino Hills, CA, USA).

| Name Vector | Description Vector | Description Fragment | Restriction sites | Description procedure | Fly Line |
| --- | --- | --- | --- | --- | --- |
| Allele switch KIWT | RIV FRT<br>MCS1 FRT<br>MCS3 pax-cherry | Schlank- | NheI; XhoI | Cutted from pGE attB KIWT | N |
| AlleleSwitch *KIWT-H215DHA* (*Sw WT*→*H215D*) | AlleleSwitch KIWT;pTV-cherry | mutagenesis in H215D site, CT HA tag | PacI, AgeI | PCR, pGE attB H215D | Y |
| AlleleSwitch *KIWT-NLS2HA* (*Sw WT*→*NLS2*) | AlleleSwitch KIWT;pTV-cherry | mutagenesis in NLS2 site; CT HA tag | PacI, AgeI | PCR, miniwhite_Schlank_NLS2-HA | Y |
| AlleleSwitch *KIWT-KIWTHA* (*Sw WT*→*WT*) | AlleleSwitch KIWT;pTV-cherry | Insertion of KIWTHA | PacI, AgeI | PCR, miniwhite_SchlankHA | N |

**S3 Table.**
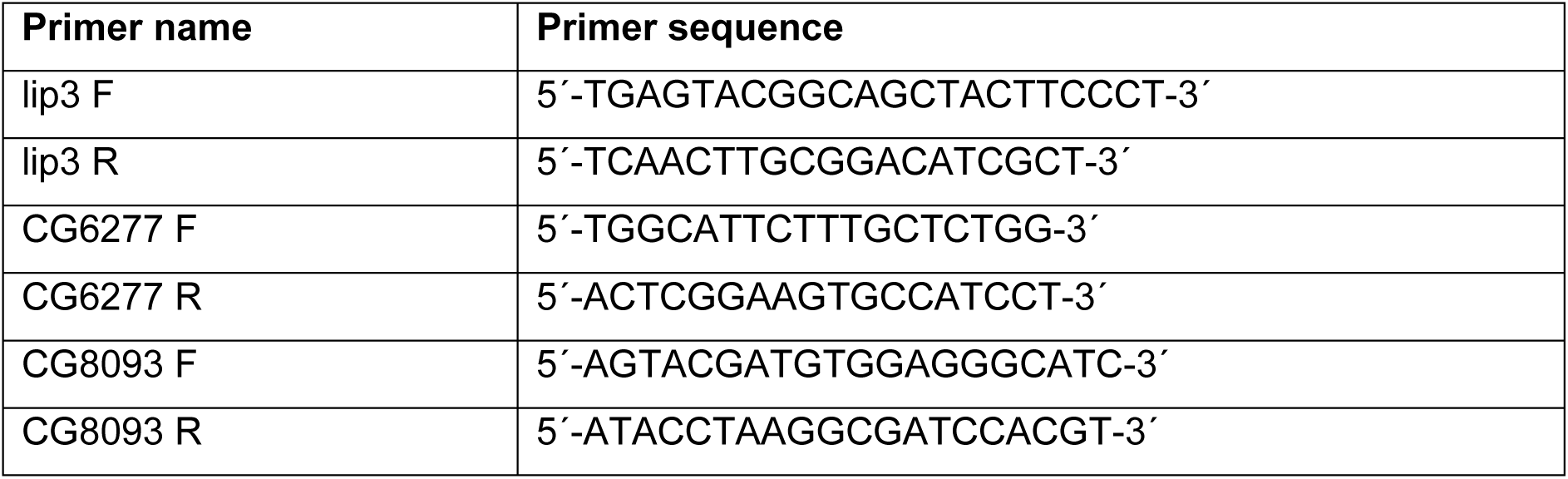
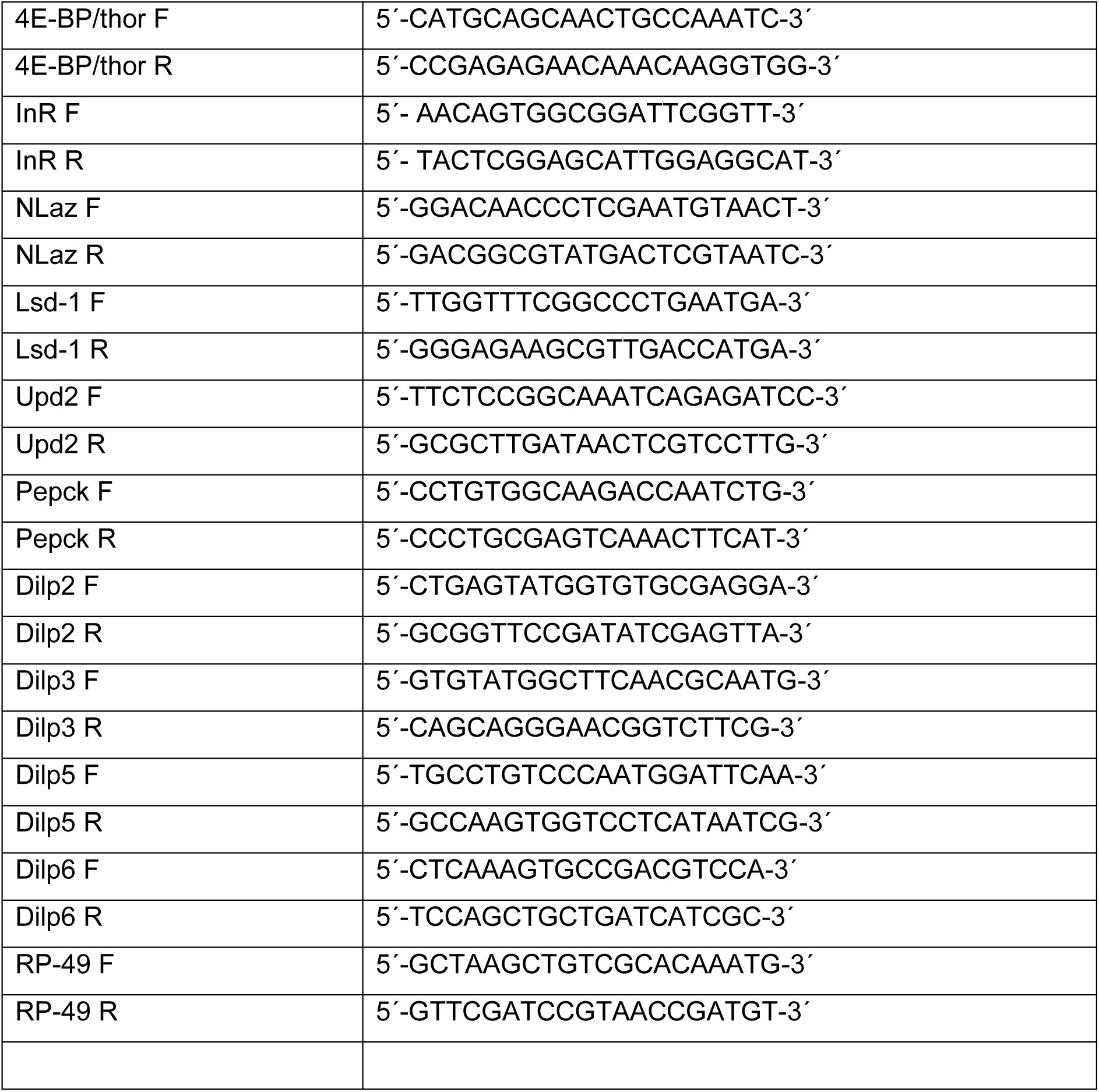
List of primers used for qRT-PCR.

**S4 Table.**
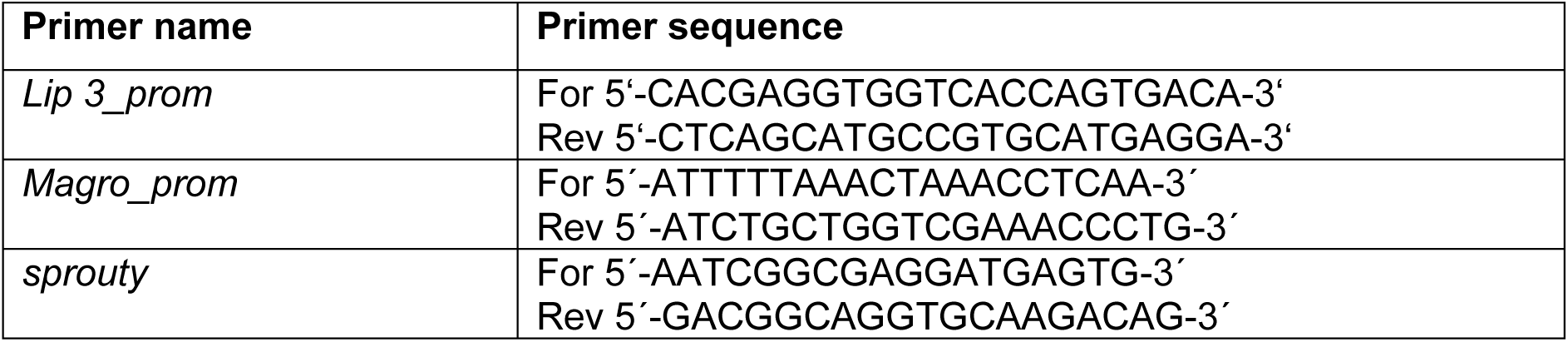
Primers used for qRT-PCR ChiP.

## Supplemental Figure Legend

**S1 Fig.**
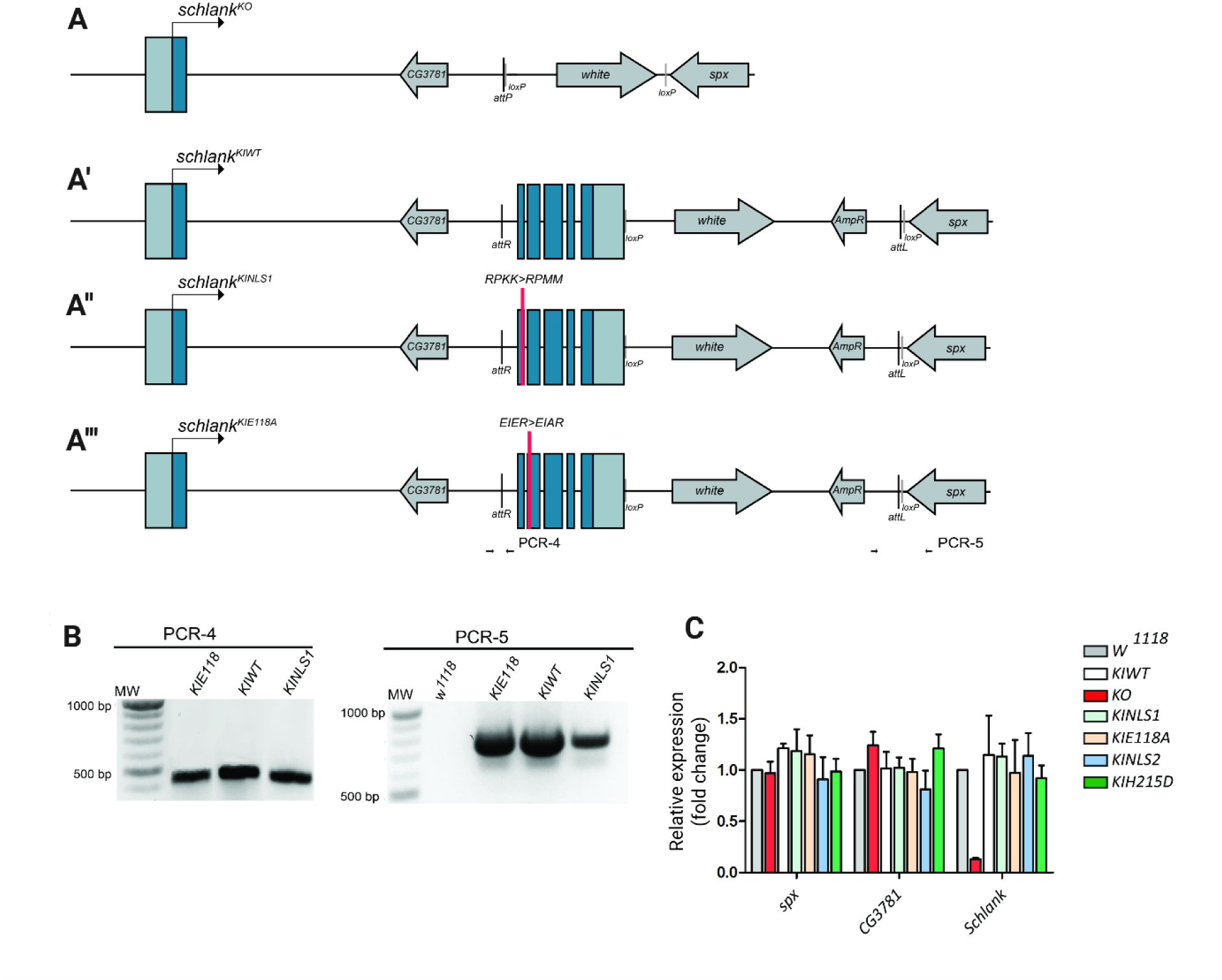
Genomic engineering of *schlank* mutant alleles. Generation and verification of *schlank* knock-in (KI) wild-type (*schlank^KIWT^*, KIWT), *schlank* knock-in NLS1 (*schlank^KINLS1^*, KINLS1), and *schlank* knock-in E118A (*schlank^KIE118A^*, KIE118A) alleles. (A) The *w^-^*, *schlank^KO^* line (Sociale et al., 2018), the founder line, was used to reintegrate target constructs into the attP site. (A’) Schematic of gDNA containing the deleted genomic DNA in a *pGE-attB* plasmid used for reintegration at the *schlank* target locus of the *schlank* founder line to generate the *schlank^KIWT^*allele. (A’’, A’’’) Schematic of gDNAs containing the deleted genomic DNA with the NLS1 and E118A mutation (depicted in red) in a *pGE-attB* plasmid used for reintegration at the *schlank* target locus. (B) PCR verification of *schlank^KIWT^*, *schlank^KINLS1^, and schlank^KIE118^*alleles. Primer pairs PCR-4 and PCR-5 (shown by → ←) used for the PCR genotyping, are designed to confirm the 5’ and 3’ attP/attB recombination events, respectively. PCR-4 used primers, which flank the attR site resulting in an amplicon of around 400 bp. The attL site is generated upon a correct reintegration event and was verified by PCR-5. The PCR is designed with one primer annealing within the ampicillin gene, while another primer anneals within the *spx* gene outside the genomic DNA region. Only the expected targeting events will yield a PCR product of about 700 bp. (C) Quantification of mRNA levels by qRT–PCR. Strongly reduced *schlank* mRNA expression in *schlank^KO^* mutants as compared with *schlank^KIWT^*, and *w^1118^* and KIWT, KIE118A, KINLS1, KIH215D (Ziegler et al., 2025), and KINLS2 (Sociale et al., 2018) animals. Residual quantities of *schlank* are due to *schlank* maternal and zygotic supply (Bauer et al., 2009). The expression of *CG3781* and *spx* is not affected *schlank^KO^* confirming the specificity of the KO. Transcripts levels of *schlank* or *of CG3781* and *spx* laying within or at the 3’ region of the *schlank* gene locus, respectively, were similar in *schlank^KIWT^* and all the engineered schlank mutant alleles and *w^1118^*confirming the expression of reintegrated gDNA under the control of the endogenous Schlank gene locus. Error bars indicate SEM.

**S2 Fig.**
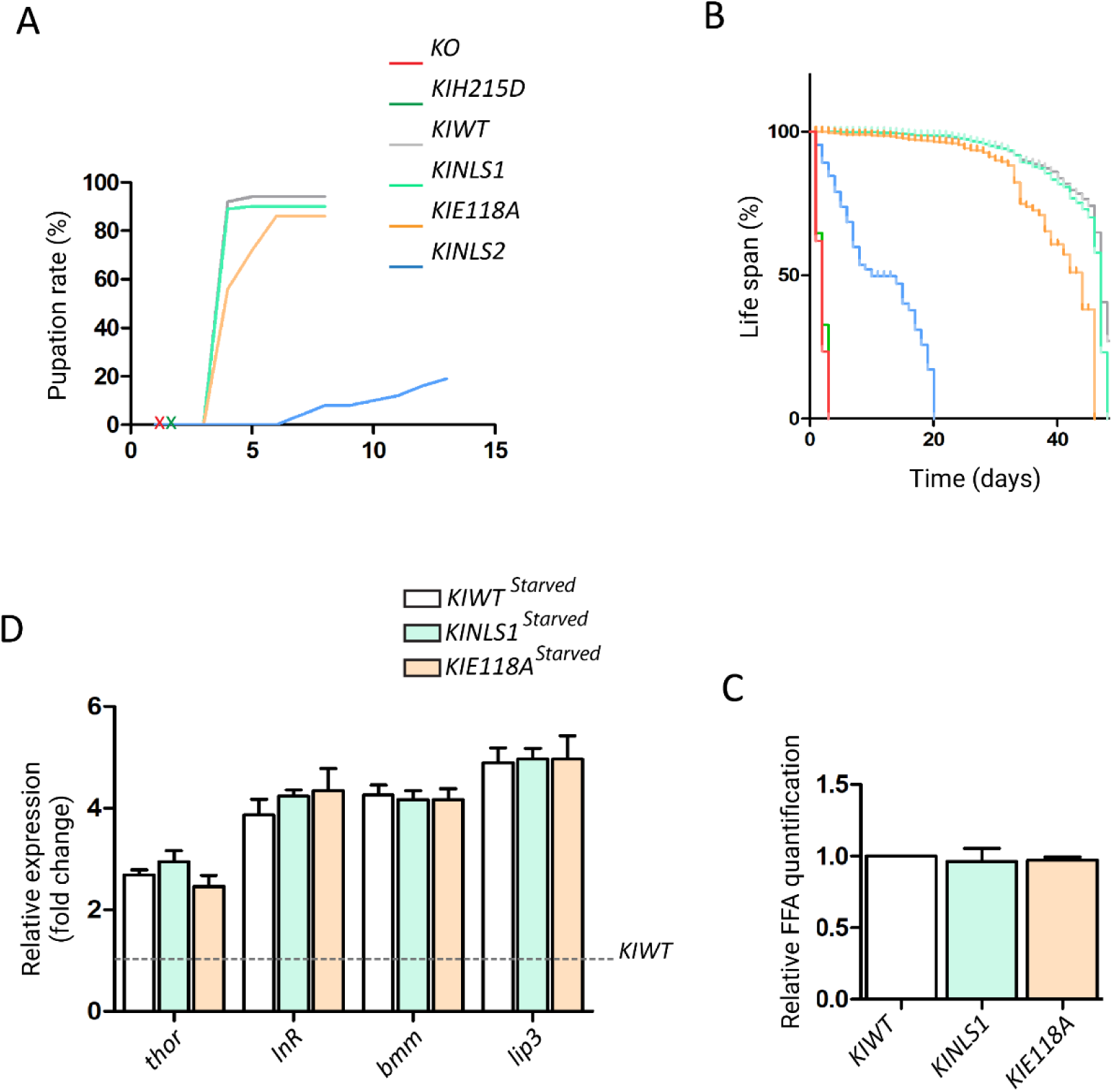
Phenotypical analysis of *KO and KIN* mutant lines. (A) Pupation rate. *KO* (red X) and *KIH215D* (green X) *animals* do not pupariate. *KINLS1* (turquois) shows the same pupation rate as WT flies, but *KIE118A KIWT* (orange) has a reduced pupation rate and a moderate developmental delay. *KINLS2* mutants show a severe developmental delay and a reduced survival rate whereby most of the surviving larvae begin to pupation at around 1day AEL. (B) Life span analysis. *KO* (red) and *KIH215D* (green) animals die approximately at 72h AEL as morphological first instar larvae. No difference in survival between *KIWT* and *KINLS1* flies (grey and turquoise), but *KIE118A* (orange) shows a moderate reduction in life span as compared to *KIWT* control animals. About 10% of *KINLS2* mutated pupae eclose as small sized adults showing severe motoric problems and most of them die within the 5 – 7 days of adulthood. (C) Starvation response is induced upon food deprivation (n=3) in KIWT, KINLS1, and KIE118A animals. Expression of genes involved in ILS (thor, InR) and known starvation induced lipases (bmm, lip3). (D) Relative free fatty acid quantification showing no significant differences among KINLS1, and KIE118A mutants in comparison with the wild-type.

**S3 Fig.**
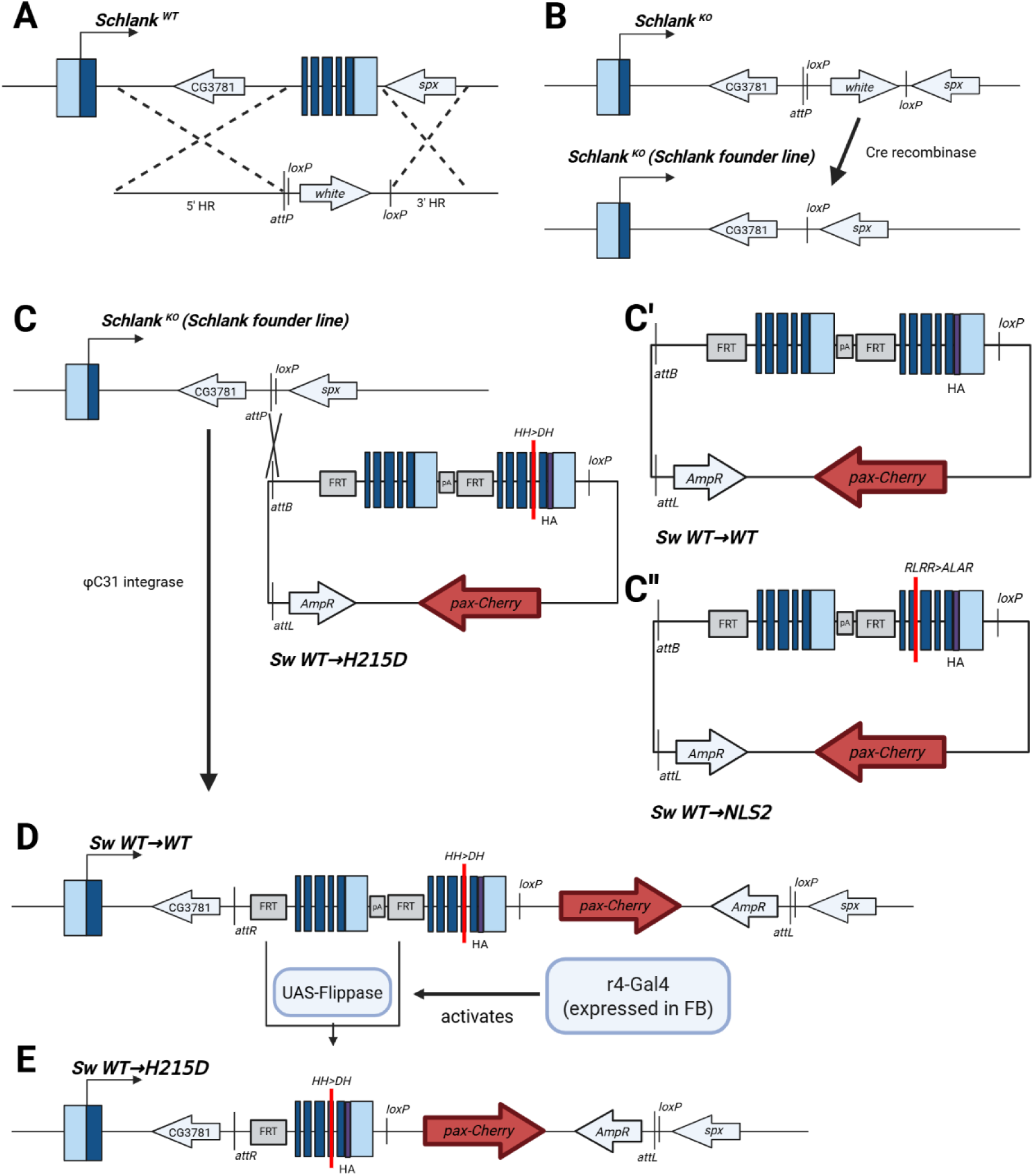
Genomic engineering of *schlank* alleles for cis-allele switching. Schematic of performed steps for the generation of Schlank cis-allele switching lines and verification of the allele switch. (A, B) *schlank^KO^* founder line and reintegration scheme of the rescuing wild type gDNA and mutant gDNA variants into the founder *schlank^KO^* founder line. The *w^-^*, *schlank^KO^* founder line was used to reintegrate target constructs in the RIVFRT MCS.pA.FRT MCS3 reintegration vector (RIV) at the attP site (Baena-Lopez et al., 2013). (C) Wild-type rescuing gDNA of schlank together with second variants of schlank gDNA, either wild type (C’) or gDNA containing a point mutation within the NLS2 site (C’’) or at H215D within the lag1p motif were integrated. (D) The genomic locus of the RIV FRT-KIWT-FRT-KIH215D line after reintegration of the target construct prior (C) or upon induction of a Flippase mediated recombination event. (E) Excision of the FRT cassette causes the locus stop expressing rescuing wild-type *schlank* and start expressing HA-tagged Schlank wild-type or NLS2 or H215D (shown as an example) variants instead.

**S4 Fig.**
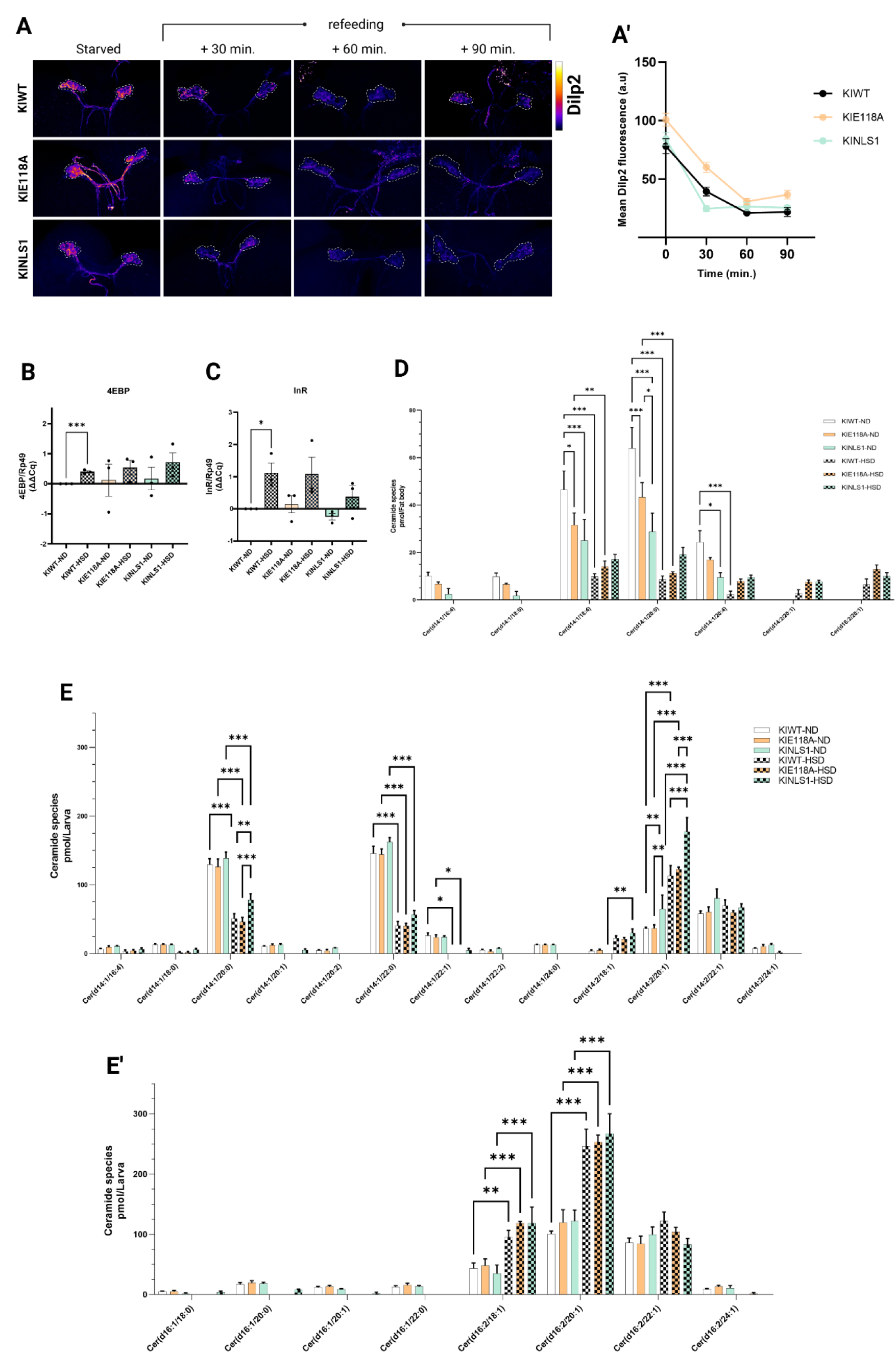
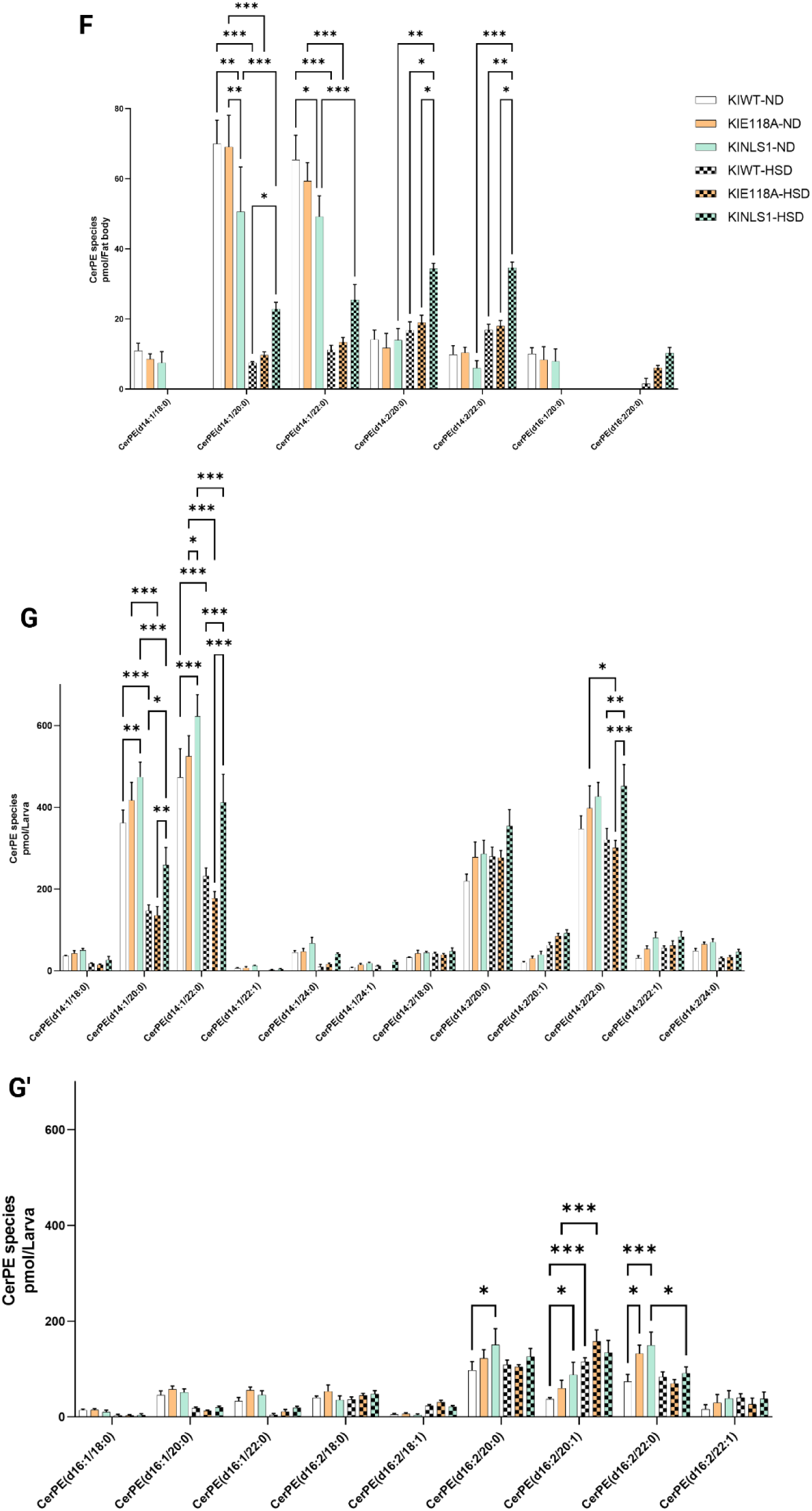
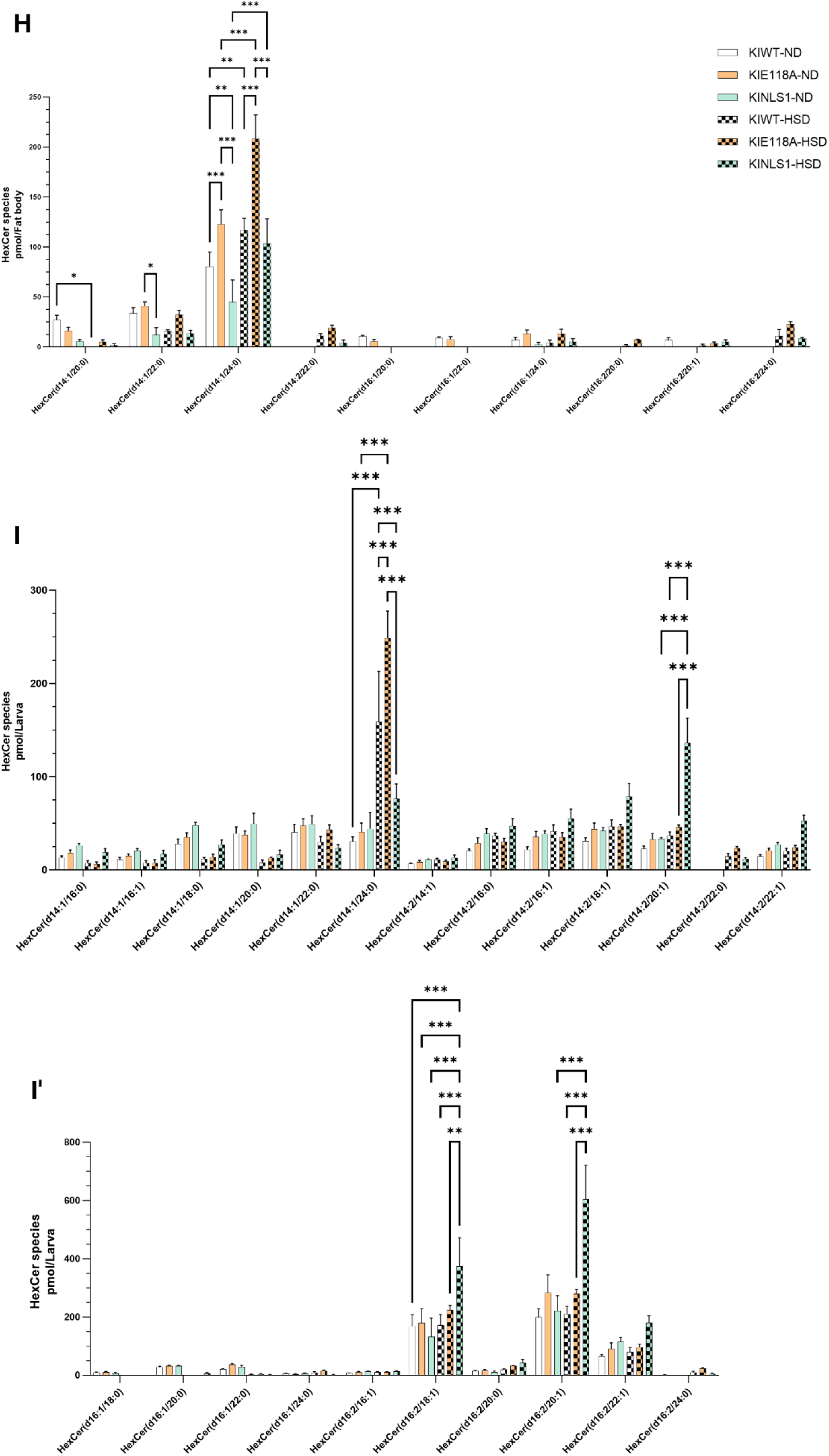
CerS E118A and NLS1 homeodomain mutants alter lipid profile, predisposing to insulin disturbance. (A) Immunostaining determining the accumulation of Dilp2 peptide through the fluorescence intensity of an α-Dilp2 antibody in IPCs of L3-instar mutated larvae (KIE118A and KINLS1) and WT (dashed outline). The animals were starved for 3 hours and immediately refed on yeast food after starvation, showing a strong accumulation of Dilp2 during starvation, but recovering the secretion after refeeding for the first 30 minutes. As time goes by (60 and 90 minutes), the secretion of Dilp2 is increased. Interestingly, the KIE118A retains Dilp2 partially over time. (A’) Quantification of relative mean Dilp2 fluorescence intensity shows Dilp2 peptide release from IPCs. Unlike the wild-type and the KINLS1, the KIE118A mutated larvae still accumulate Dilp2 when refed with yeast for 30, 60 and 90 minutes after starvation (n≥15). (B, C) Quantification of the transcript levels of FoxO target genes (B) 4ebp, and (C) InR by RT-qPCR in the whole larvae of CerS homeodomain mutants and wild-type reared on ND and HSD (n=3 biological replicates). The expression level is normalized on rp-49, and the ΔΔCq is represented and used in statistical analysis. (D) Bar chart of representative ceramide species assessed in KINLS1 and KIE118A compared with the wildtype control, and quantified relative to lipid standards in each replicate of one fat body each and shown in pmol/fat body, after rearing the animals on ND and HSD. Lipids were extracted and measured by tandem shotgun MS (n=4 biological replicates). (E, E’) Bar chart of representative ceramide species assessed in KINLS1 and KIE118A compared with the wildtype control, and quantified relative to lipid standards in each replicate of one complete larva, shown in pmol/larva, after rearing the animals on ND and HSD. Lipids were extracted and measured by tandem shotgun MS (n=4 biological replicates). (F) Bar chart of representative CerPE species assessed in KINLS1 and KIE118A compared with the wildtype control, and quantified relative to lipid standards in each replicate of one fat body each and shown in pmol/fat body, after rearing the animals on ND and HSD. Lipids were extracted and measured by tandem shotgun MS (n=4 biological replicates). (G, G’) Bar chart of representative CerPE species assessed in KINLS1 and KIE118A compared with the wildtype control, and quantified relative to lipid standards in each replicate of one complete larva, shown in pmol/larva, after rearing the animals on ND and HSD. Lipids were extracted and measured by tandem shotgun MS (n=4 biological replicates). (H) Bar chart of representative HexCer species assessed in KINLS1 and KIE118A compared with the wildtype control, and quantified relative to lipid standards in each replicate of one fat body each and shown in pmol/fat body, after rearing the animals on ND and HSD. Lipids were extracted and measured by tandem shotgun MS (n=4 biological replicates). (I, I’) Bar chart of representative HexCer species assessed in KINLS1 and KIE118A compared with the wildtype control, and quantified relative to lipid standards in each replicate of one complete larva, shown in pmol/larva, after rearing the animals on ND and HSD. Lipids were extracted and measured by tandem shotgun MS (n=4 biological replicates). Error bars from each graph mentioned above indicate ±SEM. Unpaired two-tailed t-test and one-way ANOVA followed by Tukey’s multiple comparisons tests were used to derive P-values * p<0.05, ** p<0.01, *** p<0.001.

**S5 Fig.**
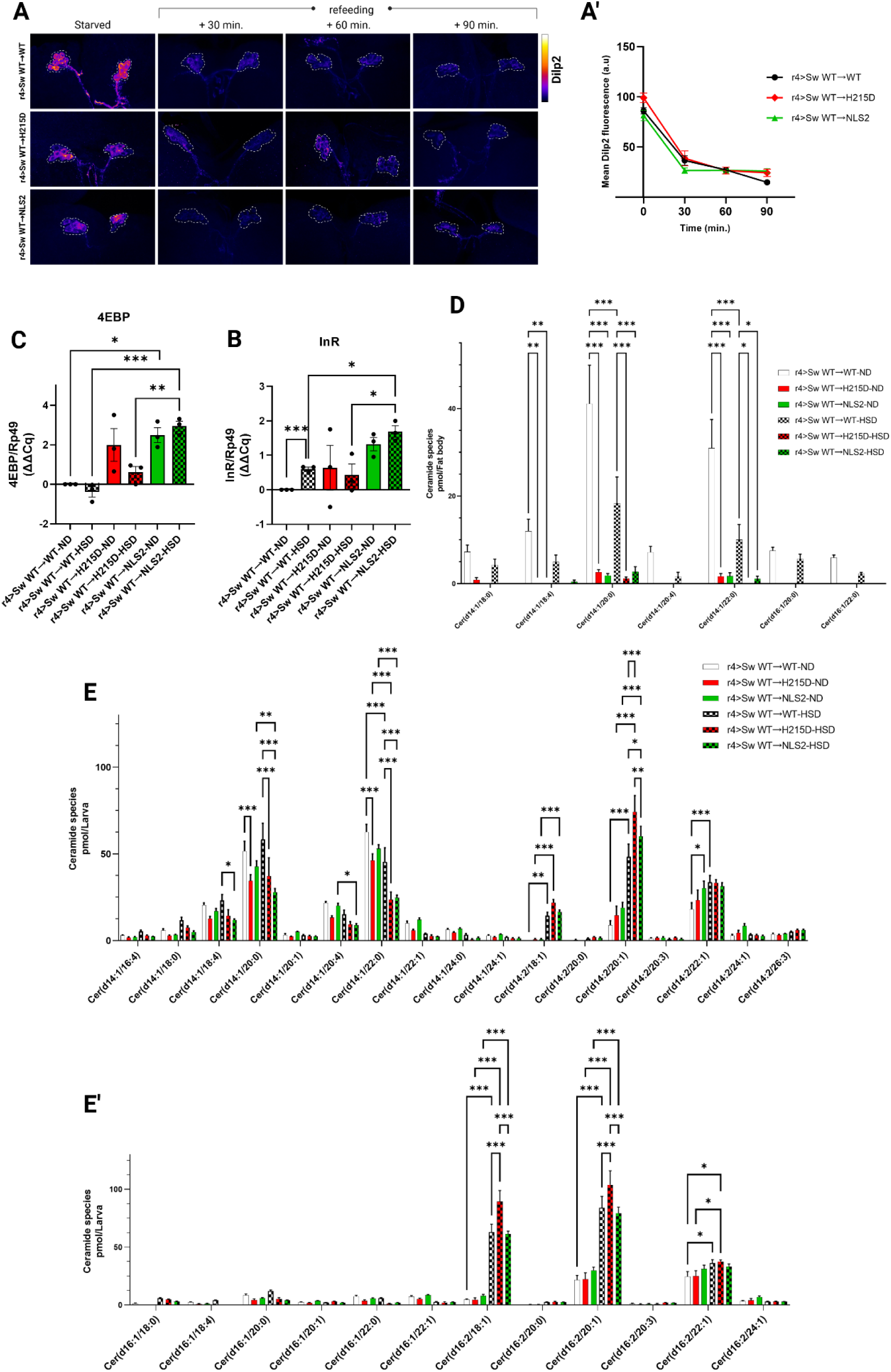
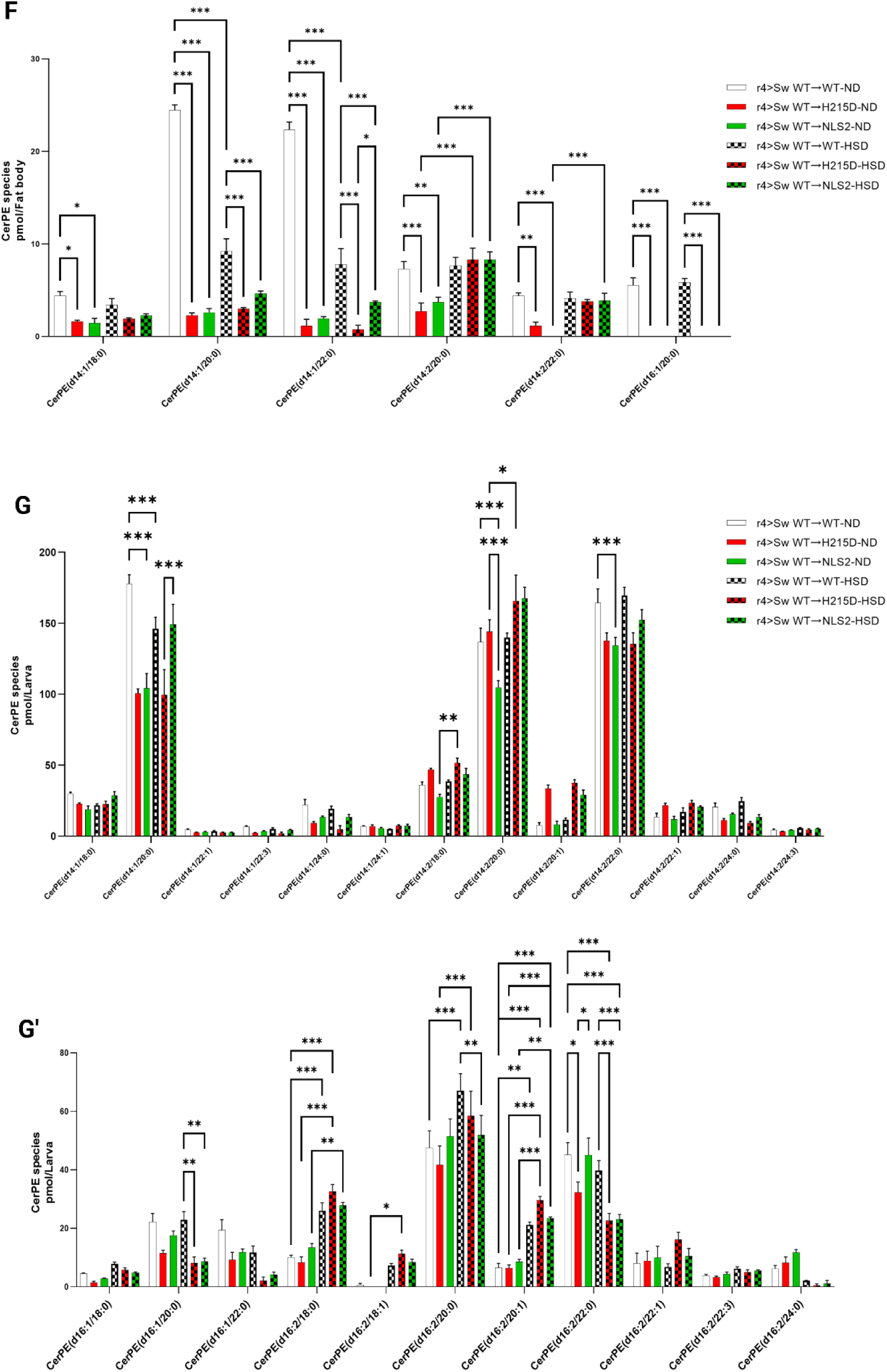
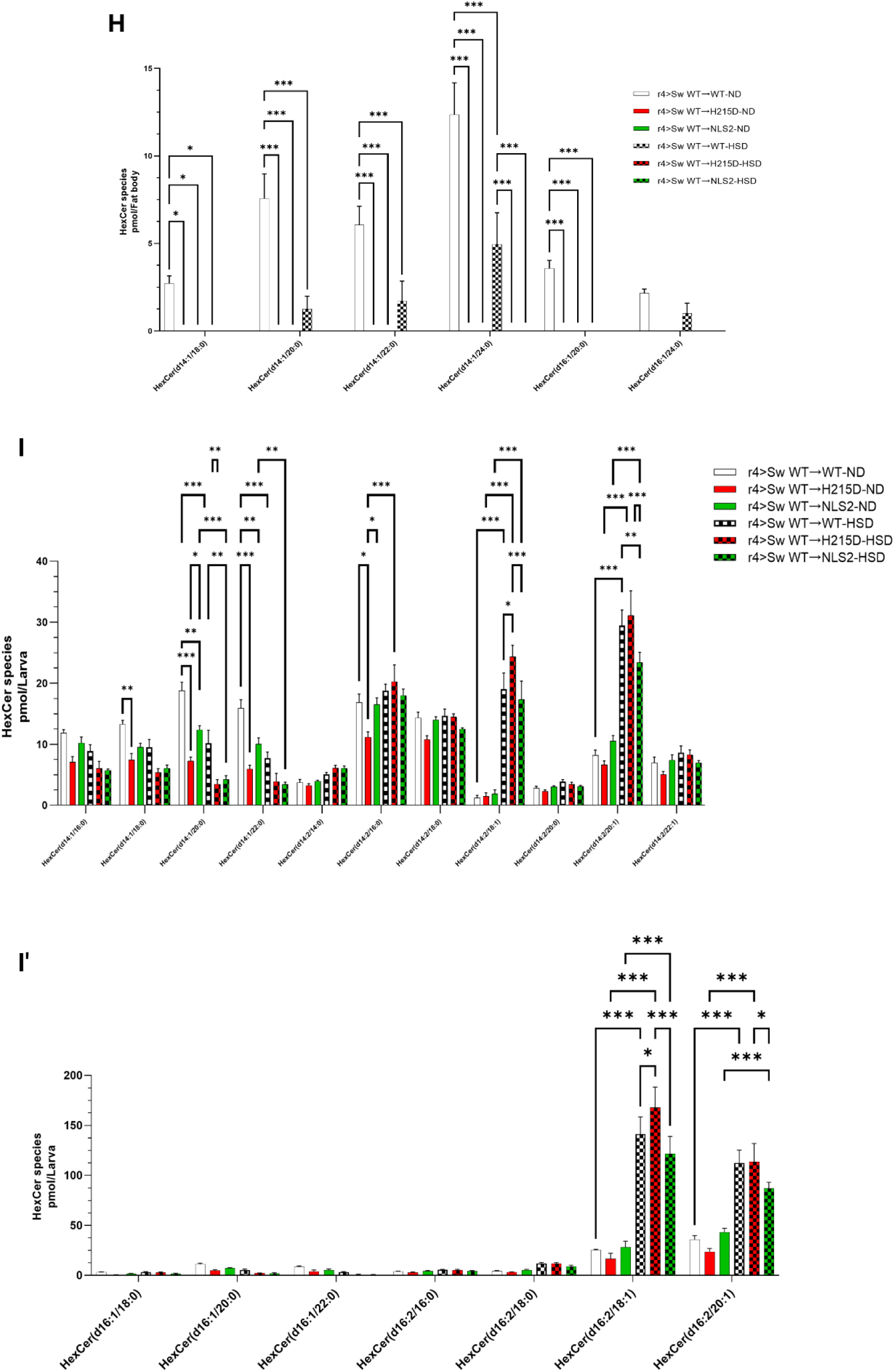
FB-expressed CerS NLS2 and H215D mutants modify lipid profile, decreasing sphingolipids. (A) Immunostaining determining the accumulation of Dilp2 peptide using an α-Dilp2 antibody in IPCs of L3-instar larvae expressing CerS mutated variants (NLS2 and H215D) and compared with the wild-type control (dashed outline). The animals were starved for 3 hours and immediately refed on yeast food after starvation, showing a strong accumulation of Dilp2 during starvation, but recovering the secretion after refeeding for the first 30 minutes. As time goes by (60 and 90 minutes), the secretion of Dilp2 is increased. No significant differences were found. (A’) Quantification of relative mean Dilp2 fluorescence intensity shows Dilp2 peptide release from IPCs (n≥15). (B, C) Quantification of the transcript levels of FoxO target genes (B) 4ebp, and (C) InR by RT-qPCR in the whole larvae of FB-expressed CerS mutants and wild-type reared on ND and HSD (n=3 biological replicates). The expression level is normalized on rp-49, and the ΔΔCq is represented and used in statistical analysis. (D) Bar chart of representative ceramide species assessed in NLS2 and H215D compared with the wildtype control, and quantified relative to lipid standards in each replicate of one fat body each and shown in pmol/fat body, after rearing the animals on ND and HSD. Lipids were extracted and measured by tandem shotgun MS (n=4 biological replicates). (E, E’) Bar chart of representative ceramide species assessed in NLS2 and H215D compared with the wildtype control, and quantified relative to lipid standards in each replicate of one complete larva, shown in pmol/larva, after rearing the animals on ND and HSD. Lipids were extracted and measured by tandem shotgun MS (n=4 biological replicates). (F) Bar chart of representative CerPE species assessed in NLS2 and H215D compared with the wildtype control, and quantified relative to lipid standards in each replicate of one fat body each and shown in pmol/fat body, after rearing the animals on ND and HSD. Lipids were extracted and measured by tandem shotgun MS (n=4 biological replicates). (G, G’) Bar chart of representative CerPE species assessed in NLS2 and H215D compared with the wildtype control, and quantified relative to lipid standards in each replicate of one complete larva, shown in pmol/larva, after rearing the animals on ND and HSD. Lipids were extracted and measured by tandem shotgun MS (n=4 biological replicates). (H) Bar chart of representative HexCer species assessed in NLS2 and H215D compared with the wildtype control, and quantified relative to lipid standards in each replicate of one fat body each and shown in pmol/fat body, after rearing the animals on ND and HSD. Lipids were extracted and measured by tandem shotgun MS (n=4 biological replicates). (I, I’) Bar chart of representative HexCer species assessed in NLS2 and H215D compared with the wildtype control, and quantified relative to lipid standards in each replicate of one complete larva, shown in pmol/larva, after rearing the animals on ND and HSD. Lipids were extracted and measured by tandem shotgun MS (n=4 biological replicates). Error bars from each graph mentioned above indicate ±SEM. Unpaired two-tailed t-test and one-way ANOVA followed by Tukey’s multiple comparisons tests were used to derive P-values * p<0.05, ** p<0.01, *** p<0.001.

## References

Alizadeh, J., da Silva Rosa, S. C., Weng, X., Jacobs, J., Lorzadeh, S., Ravandi, A., Vitorino, R., Pecic, S., Zivkovic, A., Stark, H., Shojaei, S., & Ghavami, S. (2023). Ceramides and ceramide synthases in cancer: Focus on apoptosis and autophagy. European Journal of Cell Biology, 102(3). 10.1016/j.ejcb.2023.151337

Andresini, O., Ciotti, A., Rossi, M. N., Battistelli, C., Carbone, M., & Maione, R. (2016). A cross-talk between DNA methylation and H3 lysine 9 dimethylation at the KvDMR1 region controls the induction of Cdkn1c in muscle cells. Epigenetics, 11(11), 791–803. 10.1080/15592294.2016.1230576

Azmin, M. R., Habibie, H., Filmaharani, F., Roosevelt, A., Nurhidayah, A., Pratama, M. R., Hardiyanti, W., Latada, N. P., Mudjahid, M., Yuliana, D., & Nainu, F. (2025). Aspirin-Mediated Reduction of Glucose Level and Inflammation in Drosophila melanogaster. ACS Omega. 10.1021/acsomega.4c11509

Baena-Lopez, L. A., Alexandre, C., Mitchell, A., Pasakarnis, L., & Vincent, J.-P. (2013). Accelerated homologous recombination and subsequent genome modification in *Drosophila*. Development, 140(23), 4818–4825. 10.1242/dev.100933

Bai, H., Kang, P., & Tatar, M. (2012). Drosophila insulin-like peptide-6 (dilp6) expression from fat body extends lifespan and represses secretion of Drosophila insulin-like peptide-2 from the brain. Aging Cell, 11(6), 978–985. 10.1111/acel.12000

Bandet, C. L., Tan-Chen, S., Bourron, O., Le Stunff, H., & Hajduch, E. (2019). Sphingolipid metabolism: New insight into ceramide-induced lipotoxicity in muscle cells. International Journal of Molecular Sciences, 20(3), 1–26. 10.3390/ijms20030479

Bauer, R., Voelzmann, A., Breiden, B., Schepers, U., Farwanah, H., Hahn, I., Eckardt, F., Sandhoff, K., & Hoch, M. (2009). Schlank, a member of the ceramide synthase family controls growth and body fat in Drosophila. EMBO Journal, 28(23), 3706–3716. 10.1038/emboj.2009.305

Binh, T. D., Pham, T. L. A., Men, T. T., & Kamei, K. (2019). Dysfunction of LSD-1 induces JNK signaling pathway-dependent abnormal development of thorax and apoptosis cell death in Drosophila melanogaster. Biochemical and Biophysical Research Communications, 516(2), 451–456. 10.1016/j.bbrc.2019.06.075

Brent, A. E., & Rajan, A. (2020). Insulin and Leptin/Upd2 Exert Opposing Influences on Synapse Number in Fat-Sensing Neurons. Cell Metabolism, 32(5), 786–800.e7. 10.1016/j.cmet.2020.08.017

Bülow, M. H., Wingen, C., Senyilmaz, D., Gosejacob, D., Sociale, M., Bauer, R., Schulze, H., Sandhoff, K., Teleman, A. A., Hoch, M., & Sellina, J. (2018). Unbalanced lipolysis results in lipotoxicity and mitochondrial damage in peroxisome-deficient Pex19 mutants. Molecular Biology of the Cell, 29(4), 396–407. 10.1091/mbc.E17-08-0535

Chaurasia, B., Tippetts, T. S., Mayoral Monibas, R., Liu, J., Li, Y., Wang, L., Wilkerson, J. L., Sweeney, C. R., Pereira, R. F., Sumida, D. H., Maschek, J. A., Cox, J. E., Kaddai, V., Lancaster, G. I., Siddique, M. M., Poss, A., Pearson, M., Satapati, S., Zhou, H.,…Summers, S. A. (2019). Targeting a ceramide double bond improves insulin resistance and hepatic steatosis. Science, 365(6451), 386–392. 10.1126/science.aav3722

Chavez, J. A., & Summers, S. A. (2012). A ceramide-centric view of insulin resistance. Cell Metabolism, 15(5), 585–594. 10.1016/j.cmet.2012.04.002

Chen, W. W., Chao, Y. J., Chang, W. H., Chan, J. F., & Hsu, Y. H. H. (2018). Phosphatidylglycerol Incorporates into Cardiolipin to Improve Mitochondrial Activity and Inhibits Inflammation. Scientific Reports, 8(1). 10.1038/s41598-018-23190-z

Chitkara, S., & Atilla-Gokcumen, G. E. (2025). Decoding ceramide function: how localization shapes cellular fate and how to study it. In Trends in Biochemical Sciences. Elsevier Ltd. 10.1016/j.tibs.2025.01.007

Chu, I., Chen, Y. C., Lai, R. Y., Chan, J. F., Lee, Y. H., Balazova, M., & Hsu, Y. H. H. (2022). Phosphatidylglycerol Supplementation Alters Mitochondrial Morphology and Cardiolipin Composition. Membranes, 12(4). 10.3390/membranes12040383

de María Márquez Álvarez, C., Gómez-Crisóstomo, N. P., De la Cruz-Hernández, E. N., Zazueta, C., Aguilar-Gamas, C. F., & Martínez-Abundis, E. (2023). Differential disruption on glucose and insulin metabolism in two rat models of diet-induced obesity, based on carbohydrates or lipids. Molecular and Cellular Biochemistry, 478(11), 2481–2488. 10.1007/s11010-023-04677-4

Delanoue, R., Meschi, E., Agrawal, N., Mauri, A., Tsatskis, Y., McNeill, H., & Léopold, P. (2016). Drosophila insulin release is triggered by adipose Stunted ligand to brain Methuselah receptor. Science, 353(6307), 1553–1556. 10.1126/science.aaf8430

Dobson, A. J., Ezcurra, M., Flanagan, C. E., Summerfield, A. C., Piper, M. D. W., Gems, D., & Alic, N. (2017). Nutritional Programming of Lifespan by FOXO Inhibition on Sugar-Rich Diets. Cell Reports, 18(2), 299–306. 10.1016/j.celrep.2016.12.029

Dutriaux, A., Godart, A., Brachet, A., & Silber, J. (2013). The Insulin Receptor Is Required for the Development of the Drosophila Peripheral Nervous System. PLoS ONE, 8(9). 10.1371/journal.pone.0071857

Géminard, C., Rulifson, E. J., & Léopold, P. (2009). Remote Control of Insulin Secretion by Fat Cells in Drosophila. Cell Metabolism, 10(3), 199–207. 10.1016/j.cmet.2009.08.002

Hammerschmidt, P., Steculorum, S. M., Bandet, C. L., Del Río-Martín, A., Steuernagel, L., Kohlhaas, V., Feldmann, M., Varela, L., Majcher, A., Quatorze Correia, M., Klar, R. F. U., Bauder, C. A., Kaya, E., Porniece, M., Biglari, N., Sieben, A., Horvath, T. L., Hornemann, T., Brodesser, S., & Brüning, J. C. (2023). CerS6-dependent ceramide synthesis in hypothalamic neurons promotes ER/mitochondrial stress and impairs glucose homeostasis in obese mice. Nature Communications, 14(1), 7824. 10.1038/s41467-023-42595-7

Heinitz, S., Traurig, M., Krakoff, J., Rabe, P., Stäubert, C., Kobes, S., Hanson, R. L., Stumvoll, M., Blüher, M., Bogardus, C., Baier, L., & Piaggi, P. (2024). An E115A Missense Variant in *CERS2* Is Associated With Increased Sleeping Energy Expenditure and Hepatic Insulin Resistance in American Indians. Diabetes, 73(8), 1361–1371. 10.2337/db23-0690

Herzog, R., Schwudke, D., Schuhmann, K., Sampaio, J. L., Bornstein, S. R., Schroeder, M., & Shevchenko, A. (2011). A novel informatics concept for high-throughput shotgun lipidomics based on the molecular fragmentation query language. Genome Biology, 12(1). 10.1186/gb-2011-12-1-r8

Holland, W. L., & Summers, S. A. (2008). Sphingolipids, insulin resistance, and metabolic disease: New insights from in vivo manipulation of sphingolipid metabolism. In Endocrine Reviews (Vol. 29, Issue 4, pp. 381–402). 10.1210/er.2007-0025

Hornemann, T. (2025). Sphingoid Base Diversity. In Atherosclerosis. Elsevier Ireland Ltd. 10.1016/j.atherosclerosis.2024.119091

Huang, J., Zhou, W., Dong, W., Watson, A. M., & Hong, Y. (2008). Directed, efficient, and versatile modifications of the Drosophila genome by genomic engineering. www.pnas.org/cgi/content/full/

Ingaramo, M. C., Sánchez, J. A., Perrimon, N., & Dekanty, A. (2020). Fat Body p53 Regulates Systemic Insulin Signaling and Autophagy under Nutrient Stress via Drosophila Upd2 Repression. Cell Reports, 33(4). 10.1016/j.celrep.2020.108321

Jojima, K., Edagawa, M., Sawai, M., Ohno, Y., & Kihara, A. (2020). Biosynthesis of the anti-lipid-microdomain sphingoid base 4,14-sphingadiene by the ceramide desaturase FADS3. FASEB Journal, 34(2), 3318–3335. 10.1096/fj.201902645R

Jojima, K., & Kihara, A. (2023). Metabolism of sphingadiene and characterization of the sphingadiene-producing enzyme FADS3.

Karsai, G., Lone, M., Kutalik, Z., Thomas Brenna, J., Li, H., Pan, D., von Eckardstein, A., & Hornemann, T. (2020). FADS3 is a Δ14Z sphingoid base desaturase that contributes to gender differences in the human plasma sphingolipidome. Journal of Biological Chemistry, 295(7), 1889–1897. 10.1074/jbc.AC119.011883

Khan, S. R., Ye, W. W., Van, J. A. D., Singh, I., Rabiee, Y., Rodricks, K. L., Zhang, X., Nicholson, R. J., Razani, B., Summers, S. A., Futerman, A. H., Gunderson, E. P., & Wheeler, M. B. (2025). Reduced circulating sphingolipids and CERS2 activity are linked to T2D risk and impaired insulin secretion. In Sci. Adv (Vol. 11). https://www.science.org

König, M. A., Kucharowski, N., Dancourt Ramos, D. P., Soyka, H., Wunderling, K., Bülow, T. R., Yaghmour, M. H., Thiele, C., Ache, J. M., Kuerschner, L., & Bülow, M. H. (2025). Insulin-like peptide secretion is mediated by peroxisome-Golgi interplay. 10.1101/2025.05.26.656179

Lee, S., & Dong, H. H. (2017). FoxO integration of insulin signaling with glucose and lipid metabolism. In Journal of Endocrinology (Vol. 233, Issue 2, pp. R67–R79). BioScientifica Ltd. 10.1530/JOE-17-0002

Levy, M., & Futerman, A. H. (2010). Mammalian ceramide synthases. In IUBMB Life (Vol. 62, Issue 5, pp. 347–356). 10.1002/iub.319

Liu, Y., Liao, S., Veenstra, J. A., & Nässel, D. R. (2016). Drosophila insulin-like peptide 1 (DILP1) is transiently expressed during non-feeding stages and reproductive dormancy. Scientific Reports, 6. 10.1038/srep26620

Lourido, F., Quenti, D., Salgado-Canales, D., & Tobar, N. (2021). Domeless receptor loss in fat body tissue reverts insulin resistance induced by a high-sugar diet in Drosophila melanogaster. Scientific Reports, 11(1), 3263. 10.1038/s41598-021-82944-4

Márquez Álvarez, C. de M., Gómez-Crisóstomo, N. P., De la Cruz-Hernández, E. N., El-Hafidi, M., Pedraza-Chaverri, J., Medina-Campos, O. N., & Martínez-Abundis, E. (2024). Chronic consumption of imbalance diets high in sucrose or fat induces abdominal obesity with different pattern of metabolic disturbances and lost in Langerhans cells population. Life Sciences, 336. 10.1016/j.lfs.2023.122305

McNally, B. D., Ashley, D. F., Hänschke, L., Daou, H. N., Watt, N. T., Murfitt, S. A., MacCannell, A. D. V., Whitehead, A., Bowen, T. S., Sanders, F. W. B., Vacca, M., Witte, K. K., Davies, G. R., Bauer, R., Griffin, J. L., & Roberts, L. D. (2022). Long-chain ceramides are cell non-autonomous signals linking lipotoxicity to endoplasmic reticulum stress in skeletal muscle. Nature Communications, 13(1), 1748. 10.1038/s41467-022-29363-9

Men, T. T., Binh, T. D., Yamaguchi, M., Huy, N. T., & Kamei, K. (2016). Function of lipid storage droplet 1 (Lsd1) in wing development of drosophila melanogaster. International Journal of Molecular Sciences, 17(5). 10.3390/ijms17050648

Meshrif, W. S., El Husseiny, I. M., & Elbrense, H. (2022). *Drosophila melanogaster* as a low-cost and valuable model for studying type 2 diabetes. Journal of Experimental Zoology Part A: Ecological and Integrative Physiology, 337(5), 457–466. 10.1002/jez.2580

Mullen, T. D., Hannun, Y. A., & Obeid, L. M. (2012). Ceramide synthases at the centre of sphingolipid metabolism and biology. Biochemical Journal, 441(3), 789–802. 10.1042/BJ20111626

Musselman, L. P., Fink, J. L., & Baranski, T. J. (2019). Similar effects of high-fructose and high-glucose feeding in a Drosophila model of obesity and diabetes. PLoS ONE, 14(5). 10.1371/journal.pone.0217096

Musselman, L. P., Fink, J. L., Narzinski, K., Ramachandran, P. V., Hathiramani, S. S., Cagan, R. L., & Baranski, T. J. (2011). A high-sugar diet produces obesity and insulin resistance in wild-type Drosophila. DMM Disease Models and Mechanisms, 4(6), 842–849. 10.1242/dmm.007948

Musselman, L. P., Fink, J. L., Ramachandran, P. V., Patterson, B. W., Okunade, A. L., Maier, E., Brent, M. R., Turk, J., & Baranski, T. J. (2013). Role of fat body lipogenesis in protection against the effects of caloric overload in drosophila. Journal of Biological Chemistry, 288(12), 8028–8042. 10.1074/jbc.M112.371047

Na, J., Musselman, L. P., Pendse, J., Baranski, T. J., Bodmer, R., Ocorr, K., & Cagan, R. (2013). A Drosophila Model of High Sugar Diet-Induced Cardiomyopathy. PLoS Genetics, 9(1). 10.1371/journal.pgen.1003175

Nässel, D. R., & Broeck, J. Vanden. (2016). Insulin/IGF signaling in Drosophila and other insects: Factors that regulate production, release and post-release action of the insulin-like peptides. In Cellular and Molecular Life Sciences (Vol. 73, Issue 2, pp. 271–290). Birkhauser Verlag AG. 10.1007/s00018-015-2063-3

Nässel, D. R., Kubrak, O. I., Liu, Y., Luo, J., & Lushchak, O. V. (2013). Factors that regulate insulin producing cells and their output in drosophila. Frontiers in Physiology, 4 SEP. 10.3389/fphys.2013.00252

Ni, Y. G., Wang, N., Cao, D. J., Sachan, N., Morris, D. J., Gerard, R. D., Kuro-O§, M., Rothermel, B. A., Hill, J. A., & Reynolds, D. W. (2007). FoxO transcription factors activate Akt and attenuate insulin signaling in heart by inhibiting protein phosphatases. www.pnas.org/cgi/content/full/

Nicholson, R. J., Poss, A. M., Maschek, J. A., Cox, J. E., Hopkins, P. N., Hunt, S. C., Playdon, M. C., Holland, W. L., & Summers, S. A. (2021). Characterizing a Common CERS2 Polymorphism in a Mouse Model of Metabolic Disease and in Subjects from the Utah CAD Study. The Journal of Clinical Endocrinology & Metabolism, 106(8), e3098–e3109. 10.1210/clinem/dgab155

Nirala, N. K., Rahman, M., Walls, S. M., Singh, A., Zhu, L. J., Bamba, T., Fukusaki, E., Srideshikan, S. M., Harris, G. L., Ip, Y. T., Bodmer, R., & Acharya, U. R. (2013). Survival Response to Increased Ceramide Involves Metabolic Adaptation through Novel Regulators of Glycolysis and Lipolysis. PLoS Genetics, 9(6). 10.1371/journal.pgen.1003556

P, P., Tomar, A., Madhwal, S., & Mukherjee, T. (2020). Immune Control of Animal Growth in Homeostasis and Nutritional Stress in Drosophila. Frontiers in Immunology, 11(July), 1528. 10.3389/fimmu.2020.01528

Panevska, A., Skočaj, M., Križaj, I., Maček, P., & Sepčić, K. (2019). Ceramide phosphoethanolamine, an enigmatic cellular membrane sphingolipid. In Biochimica et Biophysica Acta - Biomembranes (Vol. 1861, Issue 7, pp. 1284–1292). Elsevier B.V. 10.1016/j.bbamem.2019.05.001

Pasco, M. Y., & Léopold, P. (2012). High sugar-induced insulin resistance in Drosophila relies on the Lipocalin Neural Lazarillo. PLoS ONE, 7(5). 10.1371/journal.pone.0036583

Pascoa, T. C., Pike, A. C. W., Tautermann, C. S., Chi, G., Traub, M., Quigley, A., Chalk, R., Štefanić, S., Thamm, S., Pautsch, A., Carpenter, E. P., Schnapp, G., & Sauer, D. B. (2025). Structural basis of the mechanism and inhibition of a human ceramide synthase. Nature Structural & Molecular Biology, 32(3), 431–440. 10.1038/s41594-024-01414-3

Raichur, S., Brunner, B., Bielohuby, M., Hansen, G., Pfenninger, A., Wang, B., Bruning, J. C., Larsen, P. J., & Tennagels, N. (2019). The role of C16:0 ceramide in the development of obesity and type 2 diabetes: CerS6 inhibition as a novel therapeutic approach. Molecular Metabolism, 21,36–50. 10.1016/j.molmet.2018.12.008

Raichur, S., Wang, S. T., Chan, P. W., Li, Y., Ching, J., Chaurasia, B., Dogra, S., Öhman, M. K., Takeda, K., Sugii, S., Pewzner-Jung, Y., Futerman, A. H., & Summers, S. A. (2014). CerS2 haploinsufficiency inhibits β-oxidation and confers susceptibility to diet-induced steatohepatitis and insulin resistance. Cell Metabolism, 20(4), 687–695. 10.1016/j.cmet.2014.09.015

Rajan, A., & Perrimon, N. (2012). Drosophila cytokine unpaired 2 regulates physiological homeostasis by remotely controlling insulin secretion. Cell, 151(1), 123–137. 10.1016/j.cell.2012.08.019

Rulifson, E. J., Kim, S. K., & Nusse, R. (2002). Ablation of insulin-producing neurons in files: Growth and diabetic phenotypes. Science, 296(5570), 1118–1120. 10.1126/science.1070058

Schiffmann, S., Hartmann, D., Fuchs, S., Birod, K., Ferreirs, N., Schreiber, Y., Zivkovic, A., Geisslinger, G., Grösch, S., & Stark, H. (2012). Inhibitors of specific ceramide synthases. Biochimie, 94(2), 558–565. 10.1016/j.biochi.2011.09.007

Schleh, M. W., Caslin, H. L., Garcia, J. N., Mashayekhi, M., Srivastava, G., Bradley, A. B., & Hasty, A. H. (2023). Metaflammation in obesity and its therapeutic targeting. Science Translational Medicine, 15(723). 10.1126/scitranslmed.adf9382

Sellin, J., Wingen, C., Gosejacob, D., Senyilmaz, D., Hänschke, L., Büttner, S., Meyer, K., Bano, D., Nicotera, P., Teleman, A. A., & Bülow, M. H. (2018). Dietary rescue of lipotoxicity-induced mitochondrial damage in Peroxin19 mutants. PLoS Biology, 16(6). 10.1371/journal.pbio.2004893

Semaniuk, U., Strilbytska, O., Malinovska, K., Storey, K. B., Vaiserman, A., Lushchak, V., & Lushchak, O. (2021). Factors that regulate expression patterns of insulin-like peptides and their association with physiological and metabolic traits in Drosophila. In Insect Biochemistry and Molecular Biology (Vol. 135). Elsevier Ltd. 10.1016/j.ibmb.2021.103609

Shiffman, D., Pare, G., Oberbauer, R., Louie, J. Z., Rowland, C. M., Devlin, J. J., Mann, J. F., & McQueen, M. J. (2014). A gene variant in CERS2 is associated with rate of increase in albuminuria in patients with diabetes from ONTARGET and TRANSCEND. PLoS ONE, 9(9). 10.1371/journal.pone.0106631

Singh, A., Abhilasha, K. V., Acharya, K. R., Liu, H., Nirala, N. K., Parthibane, V., Kunduri, G., Abimannan, T., Tantalla, J., Zhu, L. J., Acharya, J. K., & Acharya, U. R. (2024). A nutrient responsive lipase mediates gut-brain communication to regulate insulin secretion in Drosophila. Nature Communications, 15(1). 10.1038/s41467-024-48851-8

Sociale, M., Wulf, A. L., Breiden, B., Klee, K., Thielisch, M., Eckardt, F., Sellin, J., Bülow, M. H., Löbbert, S., Weinstock, N., Voelzmann, A., Schultze, J., Sandhoff, K., & Bauer, R. (2018). Ceramide Synthase Schlank Is a Transcriptional Regulator Adapting Gene Expression to Energy Requirements. Cell Reports, 22(4), 967–978. 10.1016/j.celrep.2017.12.090

Spassieva, S., Seo, J. G., Jiang, J. C., Bielawski, J., Alvarez-Vasquez, F., Jazwinski, S. M., Hannun, Y. A., & Obeid, L. M. (2006). Necessary role for the Lag1p motif in (Dihydro)ceramide synthase activity. Journal of Biological Chemistry, 281(45), 33931– 33938. 10.1074/jbc.M608092200

Teleman, A. A. (2010). Molecular mechanisms of metabolic regulation by insulin in Drosophila. In Biochemical Journal (Vol. 425, Issue 1, pp. 13–26). 10.1042/BJ20091181

Turpin, S. M., Nicholls, H. T., Willmes, D. M., Mourier, A., Brodesser, S., Wunderlich, C. M., Mauer, J., Xu, E., Hammerschmidt, P., Brönneke, H. S., Trifunovic, A., Losasso, G., Wunderlich, F. T., Kornfeld, J. W., Blüher, M., Krönke, M., & Brüning, J. C. (2014). Obesity-induced CerS6-dependent C16:0 ceramide production promotes weight gain and glucose intolerance. Cell Metabolism, 20(4), 678–686. 10.1016/j.cmet.2014.08.002

Turpin-Nolan, S. M., Hammerschmidt, P., Chen, W., Jais, A., Timper, K., Awazawa, M., Brodesser, S., & Brüning, J. C. (2019). CerS1-Derived C18:0 Ceramide in Skeletal Muscle Promotes Obesity-Induced Insulin Resistance. Cell Reports, 26(1), 1–10.e7. 10.1016/j.celrep.2018.12.031

Voelzmann, A., & Bauer, R. (2010). Ceramide synthases in mammalians, worms, and insects: Emerging schemes. In Biomolecular Concepts (Vol. 1, Issues 5–6, pp. 411–422). Walter de Gruyter GmbH. 10.1515/bmc.2010.028

Voelzmann, A., Wulf, A. L., Eckardt, F., Thielisch, M., Brondolin, M., Pesch, Y. Y., Sociale, M., Bauer, R., & Hoch, M. (2016). Nuclear Drosophila CerS Schlank regulates lipid homeostasis via the homeodomain, independent of the lag1p motif. FEBS Letters, 590(7), 971–981. 10.1002/1873-3468.12125

Yu, S., Meng, S., Xiang, M., & Ma, H. (2021). Phosphoenolpyruvate carboxykinase in cell metabolism: Roles and mechanisms beyond gluconeogenesis. In Molecular Metabolism (Vol. 53). Elsevier GmbH. 10.1016/j.molmet.2021.101257

Yuan, D., Zhang, X., Yang, Y., Wei, L., Li, H., Zhao, T., Guo, M., Li, Z., Huang, Z., Wang, M., Dai, Z., Li, P., Xia, Q., Qian, W., & Cheng, D. (2024). Schlank orchestrates insect developmental transition by switching H3K27 acetylation to trimethylation in the prothoracic gland. Proceedings of the National Academy of Sciences of the United States of America, 121(35). 10.1073/pnas.2401861121

Zang, S., Wang, R., Liu, Y., Zhao, S., Su, L., Dai, X., Chen, H., Yin, Z., Zheng, L., Liu, Q., & Zhai, Y. (2024). Insulin Signaling Pathway Mediates FoxO–Pepck Axis Regulation of Glucose Homeostasis in Drosophila suzukii. International Journal of Molecular Sciences, 25(19). 10.3390/ijms251910441

Zhao, P., Huang, P., Xu, T., Xiang, X., Sun, Y., Liu, J., Yan, C., Wang, L., Gao, J., Cui, S., Wang, X., Zhan, L., Song, H., Liu, J., Song, W., & Liu, Y. (2021). Fat body Ire1 regulates lipid homeostasis through the Xbp1s-FoxO axis in Drosophila. IScience, 24(8). 10.1016/j.isci.2021.102819

Zheng, H., Yang, X., & Xi, Y. (2016). Fat body remodeling and homeostasis control in Drosophila. In Life Sciences (Vol. 167, pp. 22–31). Elsevier Inc. 10.1016/j.lfs.2016.10.019

Ziegler, A. B., Wesselmann, C., Beckschäfer, K., Wulf, A.-L., Dhiman, N., Soba, P., Thiele, C., Bauer, R., & Tavosanis, G. (2025). Lipid Disbalance Affects Neuronal Dendrite Growth and Maintenance in a Human Ceramide Synthase Disease Model. 10.1101/2024.10.31.621235

